# p53 drives premature neuronal differentiation in response to radiation-induced DNA damage during early neurogenesis

**DOI:** 10.1101/2020.06.26.171132

**Authors:** André-Claude Mbouombouo Mfossa, Mieke Verslegers, Tine Verreet, Haris bin Fida, Mohamed Mysara, Wilfred F.J. Van IJcken, Winnok H. De Vos, Lieve Moons, Sarah Baatout, Mohammed A. Benotmane, Danny Huylebroeck, Roel Quintens

**Affiliations:** Radiobiology Unit, Institute of Environment, Health and Safety, SCK•CEN, Mol, Belgium; Laboratory of Molecular Biology (Celgen), Department Development and Regeneration, KU Leuven, Leuven, Belgium; Laboratory of Neural Circuit Development and Regeneration, Department of Biology, Faculty of Science, KU Leuven, Leuven, Belgium; Center for Biomics, Erasmus University Medical Center, Rotterdam, The Netherlands; Department of Cell Biology, Erasmus University Medical Center, Rotterdam, The Netherlands; Laboratory of Cell Biology and Histology, Department of Veterinary Sciences, University of Antwerp, Antwerp, Belgium

**Keywords:** p53, microcephaly, DNA damage, radiation, premature neuronal differentiation, EMT, PMT

## Abstract

p53 regulates the cellular DNA damage response (DDR). Hyperactivation of p53 during embryonic development, however, can lead to a range of developmental defects including microcephaly. Here, we induce microcephaly by acute irradiation of mouse fetuses at the onset of neurogenesis. Besides a classical DDR culminating in massive apoptosis, we observe ectopic neurons in the subventricular zone in the brains of irradiated mice, indicative of premature neuronal differentiation. A transcriptomic study indicates that p53 activates both DDR genes and differentiation-associated genes. In line with this, mice with a targeted inactivation of *Trp53* in the dorsal forebrain, do not show this ectopic phenotype and partially restore brain size after irradiation. Irradiation furthermore induces an epithelial-to-mesenchymal transition-like process resembling the radiation-induced proneural-mesenchymal transition in glioma and glioma stem-like cells. Our results demonstrate a critical role for p53 beyond the DDR as a regulator of neural progenitor cell fate in response to DNA damage.

## Introduction

The classical function of *TP53 (Trp53* in mice) is that of a central player in the cellular response to stresses such as DNA damage through transcriptional activation of genes driving cell cycle arrest, DNA repair, senescence and apoptosis^1^. The exact outcome depends on the type and persistence of the induced damage and the level of p53 activation^2^. Collectively, this safeguards genomic stability and prevents cells from becoming malignant, a role for which p53 earned the nickname “guardian of the genome”^3^. In recent years, additional functions of p53 and its direct target genes emerged, including in autophagy, metabolism, epithelial-to-mesenchymal transition (EMT), pluripotency and differentiation, each of which can be either positively or negatively regulated by p53^4^. For this reason, p53 is now additionally considered a “guardian of cellular homeostasis”^4^.

The many biological functions and cellular processes regulated by p53 are also reflected by developmental syndromes that result from inappropriate p53 hyperactivation during embryogenesis, triggering apoptosis or restraining proliferation of certain cell types^5^. For other p53-controlled processes, like EMT and cellular differentiation, the etiology of these syndromes is currently unknown^5,6^. Many of these so-called p53-associated syndromes have been modeled in mice via introduction of mutations that affect various cellular processes converging on p53 activation. When this occurs in the brain during neurogenesis the resulting phenotypes mostly encompass microcephaly as a consequence of neuronal apoptosis^5^, which can be rescued, at least partially, by knocking out *Trp53*^7–14^. Microcephaly can also be caused by environmental factors like Zika virus (ZIKV) infection^15^ or exposure to moderate and high doses of DNA damaging ionizing radiation^16^, each at prenatal stages. The latter furthermore exemplifies the extraordinary sensitivity of the brain to DNA damage, especially during the earliest stages of embryonic neurogenesis^17^.

Premature neuronal differentiation also often underlies microcephaly because it contributes to deplete the pool of neural progenitor cells (NPCs). This can be triggered by changes in the orientation of the mitotic spindle of radial glial cells (RGCs)^18^ a type of NPCs in the embryonic neocortex. An increase of asymmetric divisions can result in the precocious generation of neurons at the expense of new RGCs, an imbalance that is often observed in primary microcephaly^19^. In contrast, microcephaly associated with early embryonic DNA damage has so far been proposed to result only from apoptosis because of the low threshold of NPCs to activate apoptosis^20,21^ as a way to maintain brain genome integrity. Whether premature neuronal differentiation can also be induced in response to embryonic DNA damage has so far not been demonstrated *in vivo*.

During neurogenesis, the differentiation of RGCs involves delamination from the apical membrane followed by radial migration towards the pial surface. This strongly resembles EMT^22^, a process during which epithelial cells lose their intercellular junctions and apical-basal polarity, reorganize their cytoskeleton and shape and reprogramme gene expression^23^. EMT is also often activated during cancer progression. An analogous mechanism underlies the proneural-mesenchymal transition (PMT) often seen in recurrent glioma with a worse prognosis, and which can be induced by radiotherapy^24,25^.

In this study, we show that irradiation during early neurogenesis leads to microcephaly as a consequence of p53-dependent apoptosis and premature neuronal differentiation. A functional genomics approach demonstrates that these two cellular outcomes depend on the regulation of both apoptosis- and differentiation-related gene signatures. The latter overlaps significantly with those activated during DNA damage-induced differentiation of mouse embryonic stem cells (ESCs). We furthermore observe molecular and cellular changes reminiscent of EMT and PMT. This indicates that the developing brain during early neurogenesis responds to ionizing radiation in a manner similar to that of gliomas and glioma-like stem cells.

## Results

### Prenatal irradiation causes DNA damage, a transient G2/M arrest, apoptosis and ultimately microcephaly

In agreement with previous observations^26^, irradiation of C57BL/6 mouse fetuses at embryonic day 11 (E11) resulted in a mild general growth deficit (male 6.8%, female 8.9%; Fig. 1a) as seen in 10-day old (P10) mice, and a strongly reduced brain weight (male 27%, female 27%; Fig. 1b), even after normalization for body weight (male 22%, female 20%; Fig. 1c). These observations were similar for male and female mice, indicating that there was no gender effect of radiation exposure on brain development.

**Figure 1.**
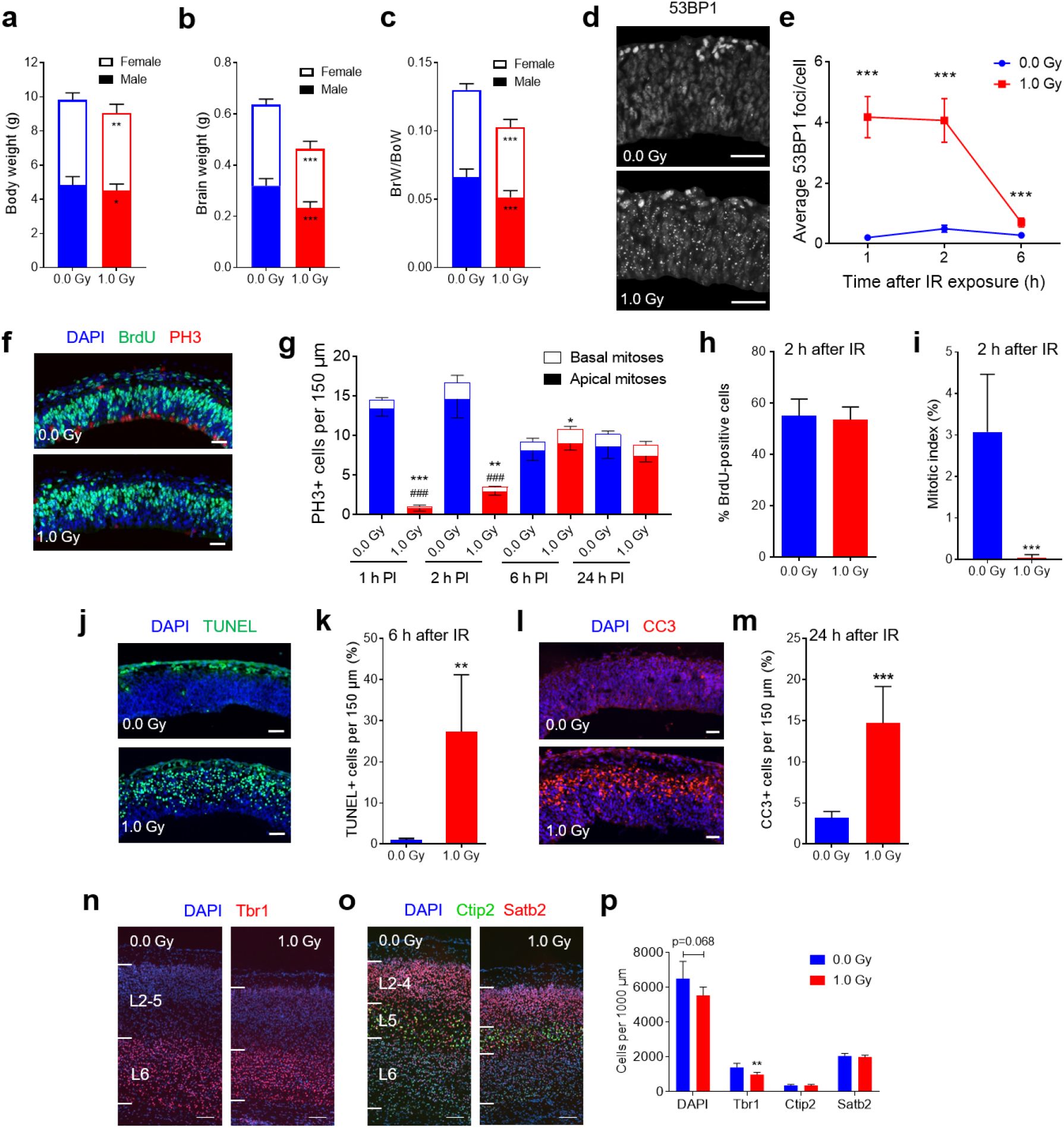
Reduced brain size and induction of DNA damage, cell cycle arrest and apoptosis in prenatally irradiated mice. (**a-c**) Body weight, brain weight and brain-to-body weight of postnatal day 10 mice after irradiation at E11. BrW: brain weight, BoW: body weight. n = 18-27. \**P* < 0.05, \*\**P* < 0.01, \*\*\**P* < 0.001 (Student’s *t*-test). (**d, e**) DNA damage was analyzed via staining for the DNA double strand break marker 53BP1. IR: Ionizing radiation. n = 3-8; \*\*\**P* < 0.001 (Student’s *t*-test); scale bar: 20 μm. (**f-i**) Staining for the late G2/mitosis marker phospho-histone 3 (PH3) and the S-phase marker BrdU indicated a transient G2/M arrest after irradiation. (**f**) Shows staining at 2 h post-irradiation. PI: post-irradiation. n = 6; \**P* < 0.05, \*\**P* < 0.01, \*\*\**P* < 0.001 (basal mitoses); ###*P* < 0.001 (apical mitoses) (Student’s *t*-test); scale bar: 20 μm. (**j, k**) Immunostaining for the apoptosis marker TUNEL indicated a strong induction of apoptosis at 6 h after irradiation. Note non-specific staining of blood vessels lining the basal membrane. n = 5-6; \*\**P* < 0.01 (Student’s *t*-test); scale bar: 20 μm. (**l, m**) Immunostaining for the apoptosis marker cleaved caspase-3 indicated a strong induction of apoptosis at 24 h after irradiation. n = 6; \*\*\**P* < 0.001 (Student’s *t*-test). (**n-p**) Immunostainings of brains at post-natal day 2 for the neocortical layer markers Tbr1 (L6), Ctip2 (L5) and Satb2 (L2-4) indicated a 30% reduction in the number of Tbr1+ L6 neurons. n = 4-6; \*\**P* < 0.01 (Student’s *t*-test); scale bar: 100 μm. In all panels data represent mean ± S.D.

Immunostaining for the DNA double-strand break (DSB) repair marker 53BP1 revealed a significant increase in DSB foci at 1 h and 2 h post-irradiation, which almost completely returned to baseline within 6 h (Fig. 1d, e). This coincided with a strong reduction in the number of phospho-histone 3 (PH3) positive apical and basal mitoses within the first 2 h (Fig. 1f, g), indicating that cell cycle arrest lasted for at least 2 h and was released after 6 h (Fig. 1g). To further investigate the cell cycle arrest, pregnant mice were injected with BrdU immediately before irradiation. A double immunostaining for incorporated BrdU and for PH3 was then performed at 2 h post-irradiation to calculate the number of cells that were irradiated while in S-phase (BrdU-positive cells, BrdU+) and progressed to mitosis (PH3+ cells). The 2-h time point was chosen because G2 and M-phase combined take around 2 h in the embryonic mouse brain^27^. Whereas the percentage of BrdU+ cells was similar in irradiated and control mice (Fig. 1h), the fraction of BrdU+/PH3+ cells over the total number of BrdU+ cells (mitotic index) showed a dramatic decrease after irradiation (Fig. 1i). This, together, indicated that cell cycle arrest was mostly due to cells arrested at the G2/M checkpoint, similar to what was found in mice irradiated at E14.5^28^.

Between 6 h and 24 h following irradiation, a strong increase in apoptosis was observed as compared to unirradiated controls, as analyzed by TUNEL (Fig. 1j, k) and cleaved caspase-3 immunostaining (Fig. 1l, m). This showed 27% and 14.8% of apoptotic cells in the neocortex at 6 h and 24 h post-irradiation, respectively. Importantly, at none of the investigated time points apoptosis was seen among the cells lining the ventricle lumen, showing that mitotic cells did not undergo apoptosis.

At one week after irradiation, the presence of apoptotic cells was no longer evident (data not shown), highlighting the transient nature of the immediate and direct effects of acute DNA damage. However, we previously showed that mice exposed to radiation at E11 have a reduced cortical thickness at this stage^29^. To identify neuronal subtypes that were lost after irradiation at E11, we performed immunostainings for markers of the different neocortical layers in brains of P2 mice. This showed that, although the patterning of the neocortex as such was not affected, the number of early-born Tbr1+ layer 6 (L6) neurons was reduced by 30% in irradiated mice as compared to non-irradiated controls (Fig. 1n, p). In contrast, the number of neurons in the more superficial layers (Ctip2, L5; Satb2, L2-4) were not affected (Fig. 1o, p). The reduction of Tbr1+ cells is consistent with loss of cells within the first day after irradiation at E11, which is around the birthdate of L6 neurons^30^. This reduction may, at least in part, be assigned to apoptosis. Together, these results confirmed the sensitivity of the brain to acute DNA damage (via radiation-induced DSBs) during the earliest stages of neurogenesis.

### Targeted *Trp53* inactivation in the embryonic dorsal forebrain partially restores DNA damage-induced microcephaly

The activation of cell cycle arrest and apoptosis following DNA damage pointed to the involvement of p53 in the early response to radiation^29,31^. Consequently, we hypothesized that genetic inactivation of *Trp53* would abrogate the negative effects of radiation on brain development. For this, we generated mice in which *Trp53* was conditionally knocked out in the neurons of the dorsal forebrain, *Emx1-Cre; Trp53*^fl/fl^ (cKO) mice (Fig. 2a), which we compared to *Trp53*^fl/fl^ (referred to as WT) littermates. Phospho-p53 staining demonstrated that loss of *Trp53* expression was lost only in the dorsal telencephalon precursors in cKO mice (Fig. 2b). Brains of unirradiated WT and cKO mice were similar at P1, suggesting that forebrain-specific removal of p53 did not affect overall brain development. After irradiation at E11, cortices of WT mice were 28% smaller than those of sham-irradiated mice (Fig. 2c-e). However, those of irradiated cKO mice were only 15% smaller (Fig. 2c-e), indicating a significant, partial rescue of the microcephalic phenotype. In WT mice irradiated at E14, cortex size was reduced with 18% (Fig. 2f), indicating reduced radiation sensitivity of E14 compared to E11 brains. This has also been reported in *Topbp1*^-/-^ mice which are more susceptible to radiation-induced apoptosis at E11 compared to E14^32^. Furthermore, at E14 when a larger fraction of cortical and hippocampal cells are *Trp53* null (Fig. S1), no difference could be observed in cortical size between irradiated cKO mice and sham-irradiated WT or cKO mice (Fig. 2f). These results suggest that while p53 seems dispensable for normal brain development, its inappropriate hyperactivation via DSBs leads to defective regulation of normal brain size.

**Figure 2:**
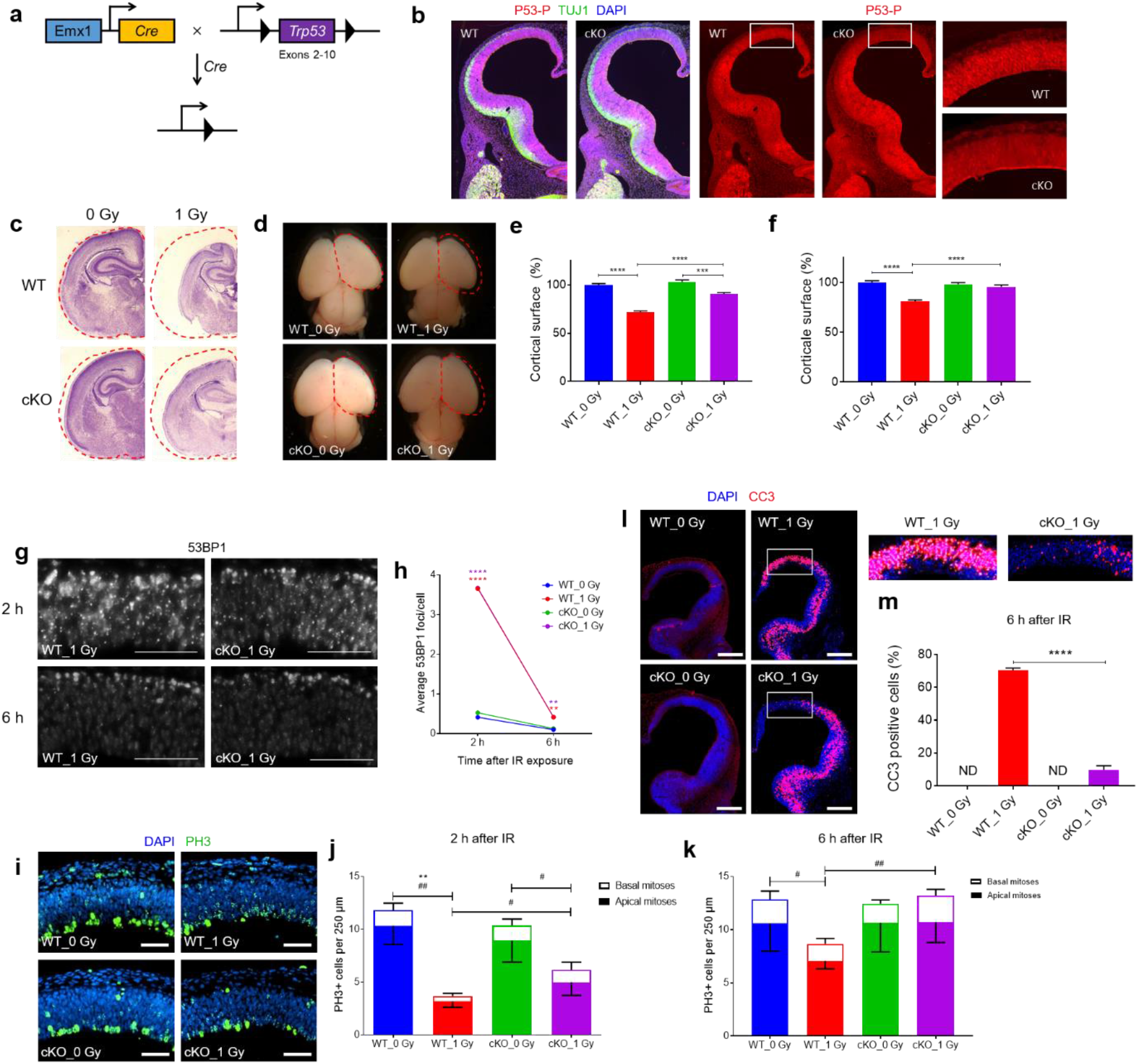
Knockout of *Trp53* in neural progenitors of the dorsal forebrain partially rescues brain size reduction after irradiation at embryonic day (E) 11. (**a**) Conditional *Trp53* knockout mice (cKO) were generated by crossing *Emx1-Cre* mice with *Trp53^fl/fl^* mice. (**b**) Staining for phosphorylated p53 (p53-P) in E11 brain at 2 h after irradiation indicates loss of p53 expression primarily in the dorsomedial pallium. Inset shows differences in p53-P staining intensity in individual cells. Please note aspecific signal in the cKO. (**c**) Representative images of haematoxylin and eosin-stained coronal sections of P1 mice of the indicated conditions after irradiation at E11. Relative brain size is indicated in red. (**d**) Representative images of dissected brains of P1 mice of the indicated conditions after irradiation at E11. Relative cerebral cortical surface is indicated in red. (**e, f**) Quantification of the relative cerebral cortical surface of P1 mice irradiated at E11 (**e**) and E14 (**f**). Two hemispheres per animal were averaged; n = 12-27 animals per condition for E11 and n = 15-20 animals per condition for E14. \*\*\**P* < 0.001; \*\*\*\**P* < 0.0001 (One-way ANOVA with correction for multiple testing according to^80^). (**g, h**) Induction of DNA damage and its initial repair are p53-independent. DNA damage was analyzed via staining for the DNA double strand break marker 53BP1. n = 6, \*\**P* < 0.01, \*\*\*\**P* < 0.0001 (One-way ANOVA with correction for multiple testing according to^80^); scale bar: 100 μm. (**i-k**) DNA damage-induced cell cycle arrest is attenuated in cKO mice. Representative images of E11 cortices at 2 h after irradiation stained for the late G2/M phase marker phospho-histone 3 (PH3) (**i**). Quantification of basal and apical mitoses per unit area at 2 h (**j**) and 6 h (**k**) after irradiation. n = 6; \*\**P* < 0.01 (basal mitoses); #*P* < 0.05, ##*P* < 0.01 (apical mitoses) (One-way ANOVA with correction for multiple testing according to^80^). (**l, m**) Induction of apoptosis is markedly reduced at 6 h after irradiation in the dorsomedial pallium of cKO mice, demonstrating the role of p53 in the regulation of DNA damage-induced apoptosis. n = 6, \*\*\*\**P* < 0.001 (One-way ANOVA with correction for multiple testing according to^80^); scale bar: 200 μm. In all panels data represent mean ± S.D.

Genetic inactivation of *Trp53* neither affected the induction nor the fast component of DSB repair (Fig. 2g, h), indicating that these are p53-independent. The induction of cell cycle arrest, however, was less efficient in cKO mice. Whereas in WT mice the numbers of apical and basal mitoses were reduced by 70% and 66%, respectively, at 2 h post-irradiation, in cKO mice the number of apical mitoses was reduced by 44%, with no significant difference in basal mitoses (Fig. 2i, j). After 6 h, the number of both basal and apical mitoses was increased in irradiated cKO mice compared to irradiated WT mice (Fig. 2k) while no difference was found in the number of mitotic cells between unirradiated and irradiated cKO mice (Fig. 2k). In agreement with p53 as a critical regulator of apoptosis, the number of apoptotic cells was drastically decreased in the dorsal telencephalon of irradiated cKO mice compared to their WT littermates at 6 h after irradiation (Fig. 2l, m). In the more lateral part of the pallium and ganglionic eminences, however, apoptosis was widespread and comparable between irradiated WT and cKO mice (Fig. 2l, m). This may at least partly explain the fact that dorsal forebrain-specific inactivation of *Trp53* only partially rescued the brain size reduction in irradiated mice.

### Whole transcriptome analysis reveals activation of a p53 signature after irradiation

To obtain more insight in the molecular mechanisms underpinning the aforementioned cellular and phenotypic changes we performed comparative genome-wide temporal RNA sequencing (RNA-seq). Changes in cortical gene expression were analyzed at 2, 6 and 12 h after irradiation of WT and cKO mice in comparison with sham-irradiated controls. In WT mice the majority of the dysregulated genes were upregulated after irradiation (Fig. 3a). Among these, 238 (after 2h), 316 (after 6 h), and 124 genes (after 12 h) were differentially expressed (adjusted *p*-value <0.05) between control and irradiated mice (Fig. S2a, S2b and Tables S1-S3). In total, 111 out of 357 (31%) differentially expressed genes overlapped between at least two time points. In contrast, only 4 out of 155 downregulated genes (2.7%) overlapped between different time points (Fig. S2b and Table S1-S3). Thus, our results showed a very dynamic transcriptional response to radiation. Compared to our previous study, in which we identified a radiation-responsive gene signature using microarrays^31^, the large majority of those genes (77 out of 83) were also identified by RNA-seq (Fig. S2c). Most of the newly identified genes changed less than 2-fold (Fig. S2d) indicating the enhanced sensitivity of RNA-seq over microarrays.

**Figure 3:**
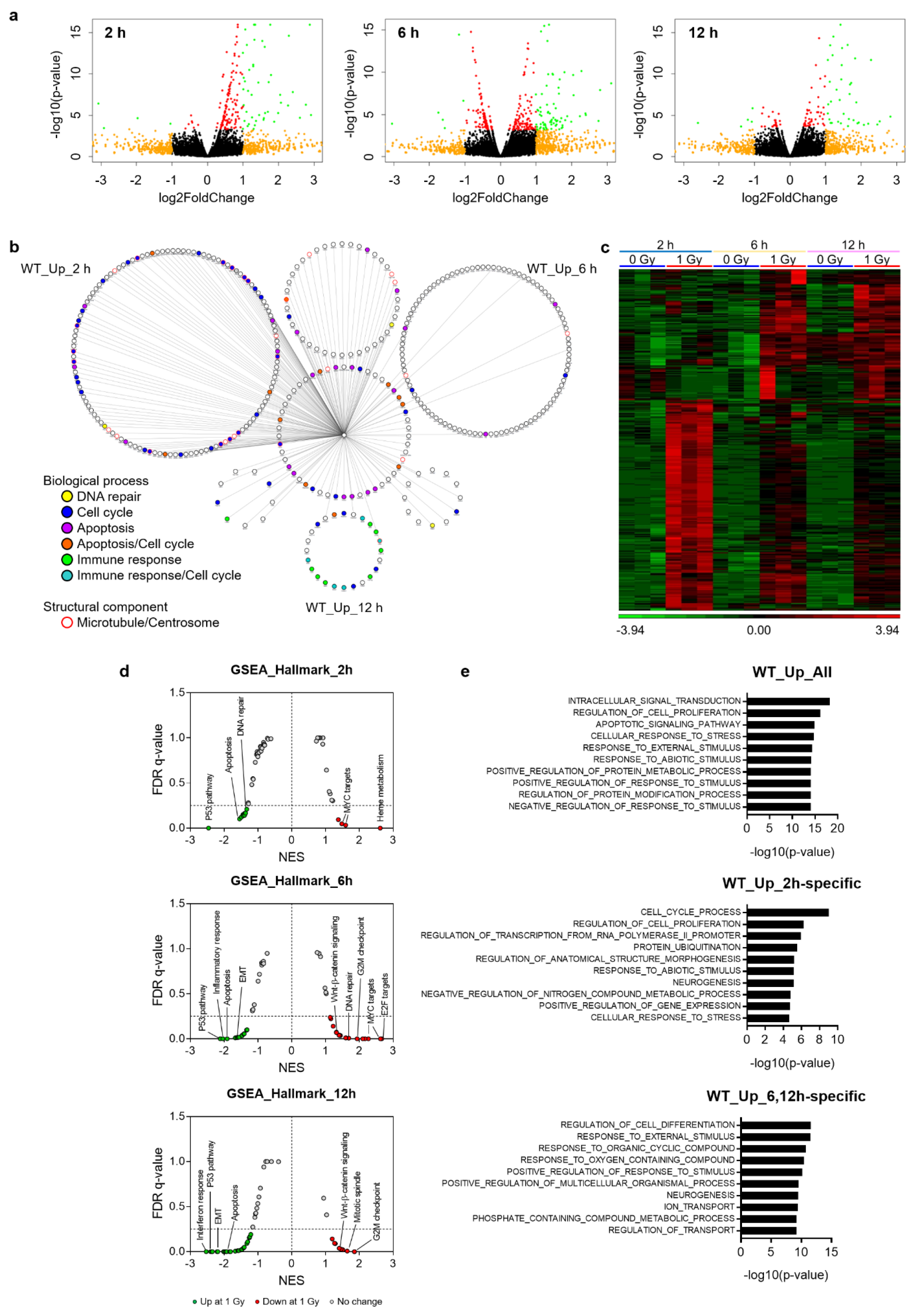
Dynamic changes in gene expression are mediated by p53. (**a**) Volcano plots representing up- and −down-regulated genes after 2 h (left), 6 h (middle) and 12 h (right). Red dots indicate genes with *P_adj_* < 0.05; orange dots indicate genes with Fold Change >|2|; green dots indicate genes with *P_adj_* < 0.05 and Fold Change >|2|. (**b**) Gene network of upregulated genes in WT mice. The central node represents p53, edges indicate direct p53 targets based on Enrichr analysis. Biological processes are indicated by node colors. (**c**) Heatmap showing unsupervised hierarchical clustering of samples based on expression levels of upregulated genes in WT mice. (**d**) Gene Set Enrichment Analysis (GSEA) for MSigDB Hallmark gene sets. Graphs depict FDR q-value versus the Nominal Enrichment Score (NES) based upon GSEA from RNA-Seq data (0 Gy versus 1 Gy). The dashed line represents the 0.25 FDR cutoff. (**e**) Gene Onology enrichment analysis was performed using the MSigDB “Investigate Gene Sets” tool. For computation of overlaps, GO Biological Processes were used as reference.

Further, and in accordance with our previous study^31^, the activated sets of genes were mostly dependent on p53 (Fig. 3b), especially after 2 h. The induction of p53-dependent genes was most pronounced (in terms of fold-change) after 2 h, and declined from 6 h onwards (Fig. S2e). Although p53-dependent genes were also activated in cKO embryonic brains (Fig. S2f), their expression was overall ~1.3-fold decreased in irradiated cKO embryos compared to WT (Fig. S2g, h). This matched with the observed ~1.3-fold reduction of Trp53 expression in cKO mice (Fig. S2h) suggesting that indeed p53 activation did not occur in the *Trp53* null fraction of cells in forebrains of cKO mice. We also investigated the effect of radiation on microRNA expression. Two hours after exposure, both increased and decreased expression of a number of microRNAs could be observed (Fig. S3a) of which miR-34a, a well-known p53 target^33,34^, was the most significant (Fig. S3b).

Gene expression profiles at 6 and 12 h were more similar to each other compared to that of 2 h post-irradiation (Fig. 3c and S2e) which was also reflected by the functional enrichment analysis. A classical p53-mediated response (cell cycle arrest, DNA repair, apoptosis) was observed after 2 h, while genes involved in the G2/M checkpoint were downregulated after 6 and 12 h (Fig. 3b, 3d-e), supporting the release of cell cycle arrest observed at 6 h. Apoptosis-related genes were induced at all time points while genes involved in the immune response were activated at later time points (Fig. 3b, 3d-e). A Gene Set Enrichment Analysis (GSEA)^35^ using curated gene sets from the MSigDB database indicated induction of genes related to radiation exposure and p53 activation at all time points (Fig. S4). The most significant overlap in this analysis was found with results from our previous study^31^, demonstrating the reproducibility of our data (Fig. S4). Interestingly, we also noticed at all time points an upregulation of genes involved in biological processes such as neurogenesis and cellular differentiation (Fig. 3e) as well as genes encoding neuronal markers (Fig. S4a, b). This coincided with a reduced expression of targets of the pluripotency regulator MYC (Fig. 3e, S4b), and other embryonic stem cell (ESC) and pluripotency gene signatures (Fig. S4). Our transcriptome analysis thus indicated that after irradiation, p53 quickly activates a classical DDR which attenuates as DNA damage repair progresses. On the other hand, a more sustained induction of genes related to neurogenesis was observed suggesting that radiation-induced neuronal differentiation may represent an additional mechanism to limit proliferation of genomically instable cells.

### Radiation exposure leads to p53-dependent premature neuronal differentiation

To investigate whether aberrant neuronal differentiation occurred in brains of irradiated fetuses, we quantified the population of RGCs (Pax6+), intermediate progenitors (IPs) (Tbr2+) and immature post-mitotic neurons (Dcx+, Tbr1+) in C57BL/6 embryos. The fraction of Pax6+ RGCs was reduced 6 h and 24 h following exposure (Fig. 4a, b). No difference was seen in the fraction of Tbr2+ IPs after 6 h (Fig. 4c, d), while ectopic Dcx+ and Tbr1+ cells could be observed in the VZ of irradiated mice (Fig. 4e-i). The cellular fate of neuronal progenitors partly depends on their plane of mitotic division, especially during the critical time window at the early stages of neurogenesis when oblique spindle orientation leads to direct neurogenesis and depletion of the progenitor pool^36^. Hence, mitotic spindle orientations were measured using double immunostaining for PH3 and the centrosomal protein *γ*-tubulin (Fig. 4j). In line with the observed increase in ectopic neurons, the fraction of horizontally dividing cells (i.e. mitotic cleavage plane <30°) was increased in irradiated brains (Fig. 4k). This, together with the fact that the number of IPs was not changed, indicated that DNA damage induces direct neurogenic divisions of RGCs.

**Figure 4:**
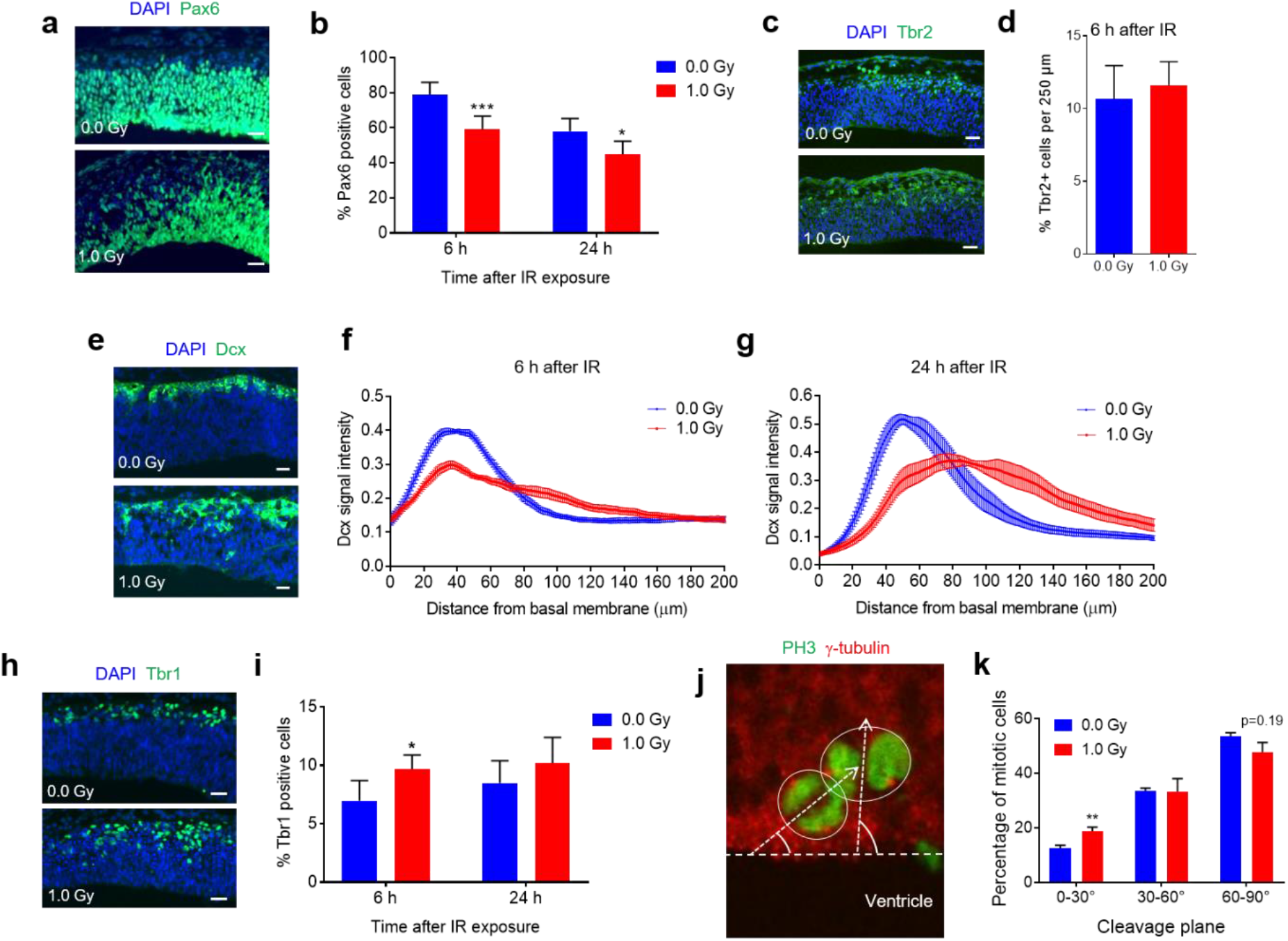
Radiation-induced DNA damage leads to premature neuronal differentiation and an increase in asymmetrically dividing radial glial cells (RGCs). (**a, b**) Reduction in the number of Pax6-positive RGCs at 6 h and 24 h after irradiation. (**c, d**) The number of Tbr2-positive intermediate progenitors was not changed after irradiation. (**e-g**) Ectopic appearance of Dcx-positive immature neurons in the ventricular zone at 6 h (**e, f**) and 24 h (**g**) after irradiation. (**h, i**) Increased number of Tbr1-positive post-mitotic pyramidal neurons at 6 h after irradiation. n = 5 (1.0 Gy, 6 h), n = 6 (all other conditions). (**j**-k) Immunostaining for the mitotic marker phospho-histone 3 (PH3) and the centrosomal protein γ-tubulin indicated an increase in the fraction of asymmetric divisions at 6 h post-irradiation. At least 150 PH3-positive cells were counted from 6 individual embryos per condition. All data represent mean + S.D. In all panels n = 6 individual embryos per condition, \**P* < 0.05, \*\**P* < 0.01, \*\*\**P* < 0.001 (Student’s *t*-test). Scale bars: 20 μm.

We then investigated whether activation of p53 regulated the process of radiation-induced premature neuronal differentiation, as suggested by our RNA-seq analysis. For this, some of the immunofluorescence experiments were repeated using *Trp53* WT and cKO mice. As in C57BL/6 mice, the fraction of Pax6+ RGCs was reduced in irradiated WT mice, both at 6 h and 24 h following irradiation (Fig. 5a-c). However, irradiated cKO mice did not show a decrease in RGCs compared to non-irradiated WT or cKO mice (Fig. 5a-c). Similarly, whereas ectopic immature Dcx+ neurons could be observed in irradiated WT mice, this was not the case in irradiated cKO littermates (Fig. 5d, e). These results indicated that radiation-induced premature neuronal differentiation was mediated via p53. To investigate whether RGCs increased their proliferation rate as a possible compensatory mechanism to account for loss of the progenitor pool, we analyzed S-phase duration and total cell cycle length using a cumulative EdU/BrdU pulse labeling approach (Fig. 5f). This indicated that, at least at 24 h after irradiation, neither the length of the cell cycle nor of the S-phase were affected (Fig. 5g, S5).

**Figure 5:**
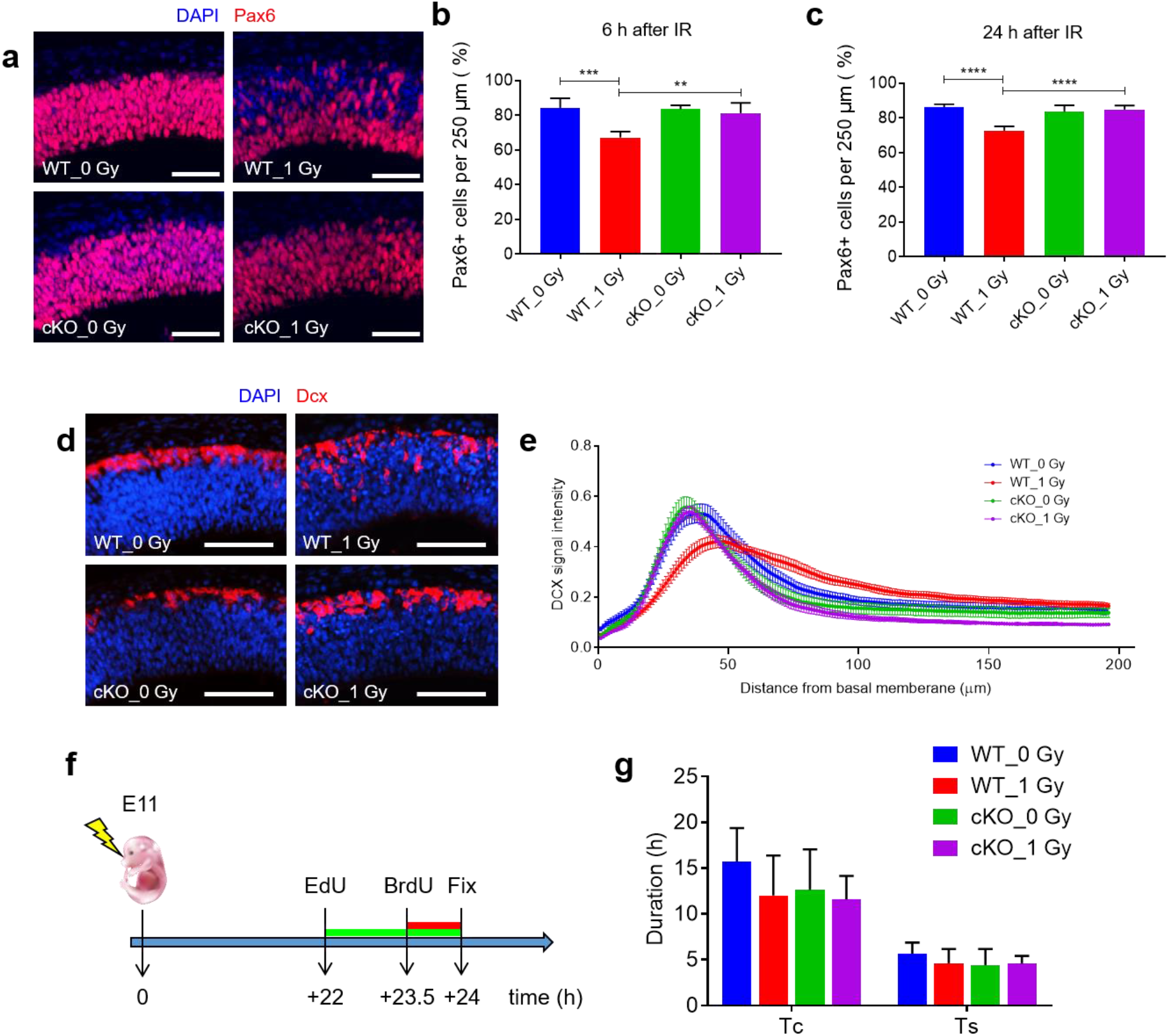
Ablation of *Trp53* prevents premature neuronal differentiation in irradiated fetuses. (**a-c**) Loss of Pax6 positive radial glia cells (RGCs) is rescued in cKO mice. Representative images of E11 cortices at 6 h after irradiation stained for the RGC marker Pax6 (**a**). Quantification of Pax6 positive cells per unit area at 6 h (**b**) and 24 h (**c**) after irradiation. n = 6; \*\**P* < 0.01, \*\*\**P* < 0.001, \*\*\*\**P* < 0.0001 (One-way ANOVA with correction for multiple testing according to^80^). (**d, e**) Radiation-induced premature neurogenesis is absent in cKO mice. Representative images of E11 cortices at 6 h after irradiation stained for the immature neuronal marker Dcx (**d**). Quantification of Dcx fluorescence intensity along the neocortex at 6 h after irradiation (**e**). n = 6 (WT_0 Gy) or n = 5 (all other conditions); scale bar: 100 μm. (**f**) Graphical representation of the sequential EdU/BrdU administration paradigm. (**g**) Quantification of total cell cycle (Tc) and S-phase (Ts) duration, based on EdU and BrdU incorporation. In all panels data represent mean + S.D. Mouse image courtesy of DataBase Center for Life Science.

A potential role of p53 in radiation-induced neuronal differentiation had previously also been suggested based on *in vitro* experiments^37,38^. We confirmed these results, and showed that irradiation of primary mouse NPCs (with 1 Gy of X-rays) diminished proliferation and induced differentiation from bipolar cells into multipolar neurons (Fig. S6a). Likewise, irradiation of Neuro-2a neuroblastoma cells (with 8 Gy of X-rays) arrested their proliferation and augmented multipolar neurite outgrowth, thereby resembling differentiated neurons, rather than bipolar NPC-like cells that were observed when Neuro-2a cells differentiated in the presence of retinoic acid (Fig. S6b). Pre-treatment of Neuro-2a cells with the p53 inhibitor alpha-pifithrin (a-PFT), reduced neurite outgrowth indicating that this is p53-dependent (Fig. S6c). Together, our results strongly suggested that p53 status not only influences DNA damage-induced cell cycle arrest and apoptosis, but also premature neuronal differentiation. We therefore propose that p53 activation is responsible for premature differentiation of NPCs in response to acute radiation-induced DNA damage.

### Convergence and divergence of gene expression changes in prenatally irradiated and *Magoh*^+/-^ microcephalic mice

Gene expression changes in irradiated mouse brains overlap significantly^31^ with those observed in the genetic microcephaly mouse model *Magoh*^+/-^ ^39^. This suggests that the converging phenotypic features (i.e. apoptosis, premature neuronal differentiation and microcephaly) directly originate from these transcriptional changes. However, in a follow-up study of the *Magoh*^+/-^ mouse model it was shown that prolonged mitosis, as is seen in *Magoh*^+/-^ NPCs induces p53-dependent apoptosis, while premature neuronal differentiation was found to be p53-independent^40^. We therefore hypothesized that radiation induced a specific p53-dependent gene signature responsible for premature neuronal differentiation.

Thus, we re-evaluated here the previously observed overlap in gene expression changes between prenatally irradiated brains and embryonic brains of *Magoh*^+/-^ mice^31^, now distinguishing between genes that were commonly upregulated in irradiated and *Magoh*^+/-^ mice (*Overlap*), genes that were only upregulated in *Magoh*^+/-^ mice (*Magoh_unique*) and genes that were only upregulated in irradiated mice (*IR_unique*) (Fig. 6a). Transcription factor enrichment analysis showed that *Overlap* genes are mainly regulated by p53 (Fig. 6b) and involved in typical p53-mediated pathways such as DNA damage response, apoptosis and cell cycle checkpoints (Fig. 6c and Table S4). Also the *IR_unique* genes were highly enriched among p53 targets (Fig. 6d). However, these correlated not only with classical p53-related functions but also forebrain development and neurogenesis (Fig. 6e and Table S5). In contrast, *Magoh_unique* genes, were enriched in targets of erythroid differentiation factors like EKLF, GATA1 and TAL1 as well as critical pluripotency factors like MYC and OCT-4 (Fig. 6f), and in biological functions related to ribosome biogenesis and cell proliferation (Fig. 6g and Table S6). Of note, MYC targets were downregulated in the brains of irradiated mice (Fig. S3).

**Figure 6:**
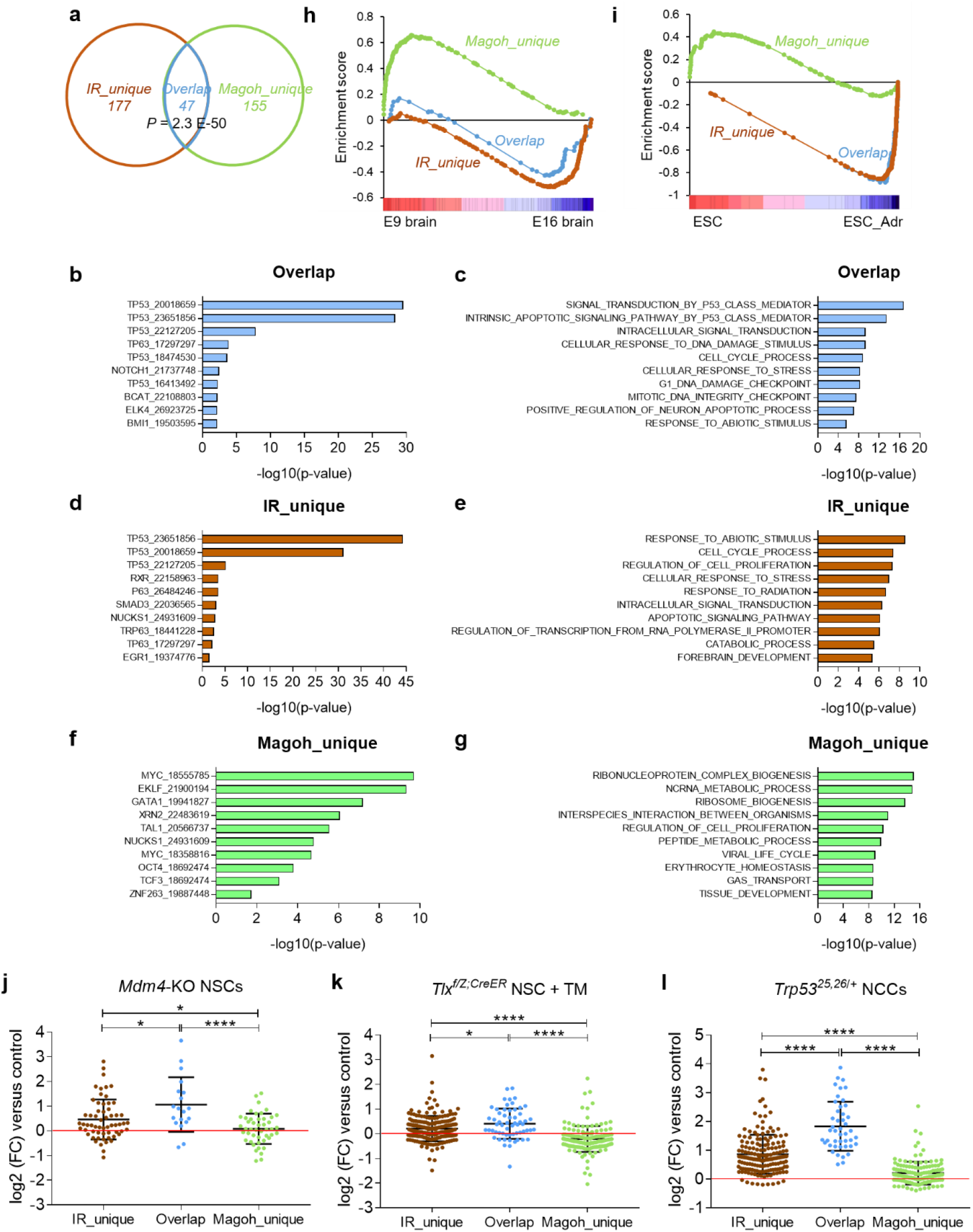
Radiation-induced DNA damage specifically regulates p53-dependent genes with possible functions in brain development and stem cell differentiation. (**a**) Venn diagram representing overlapping gene expression profiles between brains of E11 mouse fetuses at 2 h after irradiation and E10.5 *Magoh*^+/-^ mice. *IR_unique* genes are uniquely upregulated in irradiated mice, *Overlap* genes are upregulated both in irradiated and *Magoh*^+/-^ mice, *Magoh_unique* genes are uniquely upregulated in *Magoh*^+/-^ mice. (**b**) Enrichment of predicted regulating transcription factors for *Overlap* genes. Numbers indicate PubMed IDs for the publications related to the respective gene sets. (**c**) Gene Ontology enrichment for *Overlap* genes was analyzed using the *Investigate Gene Sets* tool from the MSigDB. (**d**) Enrichment of predicted regulating transcription factors for *IR-unique* genes, performed as in (**b**). (**e**) Gene Ontology enrichment for *IR_unique* genes performed as in (**c**). (**f**) Enrichment of predicted regulating transcription factors for *IR-unique* genes, performed as in (**b**). (**g**) Gene Ontology enrichment for *IR_unique* genes performed as in (**c**). (**h**) Gene Set Enrichment Analysis (GSEA) of *IR_unique, Magoh_unique* and *Overlap* genes between mouse brains at E9 and E16. *IR_unique* (FDR *q* < 0.001) and *Overlap* genes (FDR *q* = 0.10) are enriched in E16 brains compared to E9 brains. In contrast, *Magoh_unique* genes (FDR *q* < 0.001) are enriched in E9 brains. Microarray data from embryonic brain gene expression are from E-MTAB-2622 (ArrayExpress). (**i**) GSEA of *IR_unique, Magoh_unique* and *Overlap* genes in mouse R1E embryonic stem cells (ESCs) treated with the DNA damage inducing agent Adriamycin (Adr) which triggers their differentiation^75^. *IR_unique* (FDR *q* < 0.001) and *Overlap* genes (FDR *q* < 0.001) are enriched in Adr-treated ESCs, while *Magoh_unique* genes (FDR *q* < 0.002) are enriched in untreated ESCs. Microarray data from adriamycin-treated ESCs are from GSE26360 (Gene Expression Omnibus). (**j-l**) Gene expression changes of *IR_unique, Magoh_unique* and *Overlap* genes in case studies of *Mdm4* knockout neural stem cells (*Mdm4*-KO NSCs)^42^, *Tlx*-deficient NSCs (*Tlx^f/Z;CreER^* NSC + TM)^43^, and neural crest cells constitutively expressing moderate levels of p53 (*Trp53*^25,26/+^ NCCs)^45^. Data represent mean ± S.D. \**P* < 0.05, \*\*\*\**P* < 0.0001 (One-way ANOVA with Tukey’s correction for multiple comparisons).

To further discern differences between these three gene sets, we evaluated their expression profiles during normal embryonic brain development and in mouse ESCs undergoing DNA damage-induced differentiation. In both these cases, *Magoh_unique* genes and radiation-induced genes (*Overlap* + *IR_unique* genes) showed completely opposite expression profiles. GSEA analysis demonstrated that *Magoh_unique* genes are extensively downregulated (FDR *q* <0.001) during brain development. In contrast, *Overlap* (FDR *q* = 0.01) and especially *IR_unique* genes (FDR *q* <0.001) showed a strong upregulation (Fig. 6h), which is typical for genes involved in neuron differentiation^31^. Also in mouse ESCs undergoing DNA-damage induced differentiation we observed a strong upregulation of radiation-induced genes while *Magoh_unique* genes were mostly downregulated (Fig. 6i). In line with this, it was recently shown that p53 regulates the elongated G1 phase of pluripotent ground state ESCs, which is lost during culture in serum-supplemented conditions or by inactivating p53^41^. Evaluation of the expression patterns of the three gene signatures in WT versus *TP53*^-/-^ ground state and serum ESCs^41^ revealed that *Overlap* genes were strongly downregulated by *TP53* knockout in both cell lines (Fig. S7c, d, g), while *Magoh_Unique* genes were mostly affected by the cell culture conditions (Fig. S7e, f, g). *IR_Unique* genes, on the other hand, were both significantly reduced by loss of P53 and highly changed by serum supplementation (Fig. S7a, b, g). The fact that the effect of the *TP53* knockout on these genes was more pronounced in ground state ESCs argues for a potential role in nervous system development as this pathway was especially affected in these cells^41^.

Furthermore, our GSEA analysis (Fig. S4) indicated large similarities between the radiation-induced gene expression profile and that of other relevant experimental models. These include the neuroepithelium of knockout mice for the negative p53 regulator *Mdm4*^42^ and neural stem cells (NSCs) deficient for the neurogenesis regulator *Tlx*^43^ which interacts with the p53 pathway to control postnatal NSC activation^44^. Intersectional analysis showed indeed that *Overlap* and *IR_unique* genes were in general upregulated in these conditions, whereas *Magoh_unique* genes were not (Fig. 6j, k). Also, a recent study investigated time-dependent gene expression responses in neural crest cells displaying constitutively moderate p53 activation^45^. Again, we observed a substantially higher overlap with this model between *Overlap* and *IR_unique* genes, than with *Magoh_unique* genes (Fig. 6l). Altogether, these results show that both convergent and divergent gene expression changes occur in the brains of irradiated and those of *Magoh*^+/-^ mouse embryos. This may explain the phenotypic similarities (p53-dependent apoptosis) and differences (p53-dependent neuronal differentiation) between these models.

### Radiation induces an EMT-like mechanism in the embryonic brain reminiscent of the radiation-induced PMT in GSCs

Another important developmental pathway that was affected in the brains of irradiated mice was EMT (Figs. 3d, S4c). RNA-seq showed a time-dependent upregulation of EMT hallmark genes and mesenchymal markers such as *Acta2, Cthrc1, Exoc4, Lum, Myl9, Serpine2, Spp1*, and *Tagln* in brains of irradiated mice, especially at the later time points (Fig. 7a). Delamination of RGCs from the apical membrane to allow radial migration of post-mitotic cells to the basal membrane resembles EMT^22^. This coincides with changes in expression of AJ proteins and disruption of the apical AJ belt^46,47^ as is also observed when embryonic mouse brains are infected with the non-structural protein NS2A of the Zika Virus (ZIKV-NS2A)^48^. This impairs cortical neurogenesis through premature differentiation of RGCs by disrupting AJ formation and reduced expression of AJ components such as ZO-1 and β-catenin^48^.

**Figure 7:**
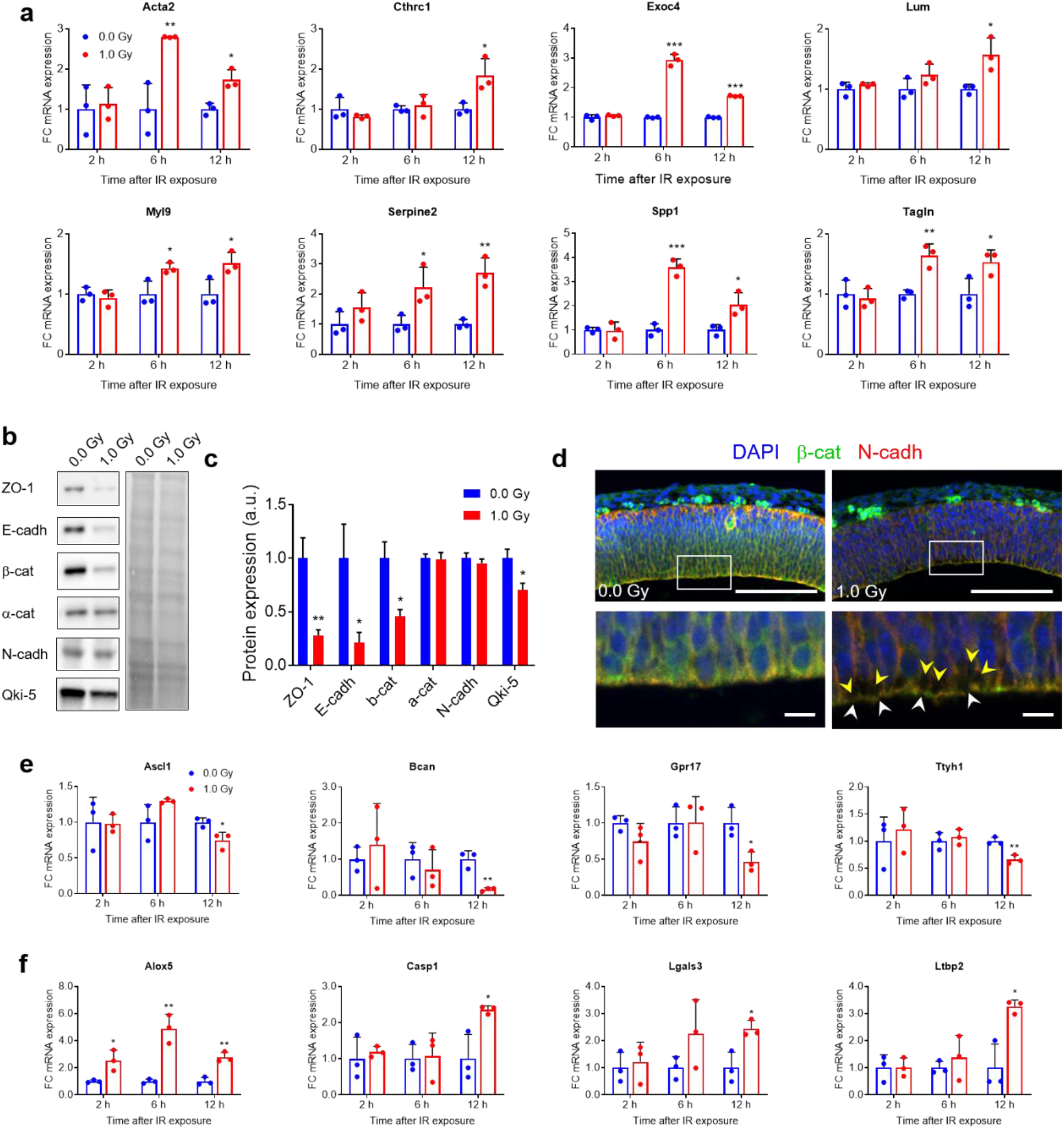
Time-dependent induction of epithelial-to-mesenchymal transition (EMT)-related genes and induction of an EMT-like process after irradiation. (**a**) Gene expression of EMT-related genes. Data represent fold changes of mRNA expression relative to 0.0 Gy for every time point (normalized to mean of 1). Data represent mean + S.D. \**P* < 0.05; \*\**P* < 0.01; \*\*\**P* < 0.001 (Student’s *t*-test with correction for multiple comparisons according to^80^). (**b**) Western blotting at 6 h after irradiation for adherens junction (AJ) complex proteins (left) were normalised for total protein (right). (**c**) Semi-quantitative analysis of the intensities of the western blot results. n = 6; \**P* < 0.05, \*\**P* < 0.01 (Student’s *t*-test). (**d**) Representative images of immunostaining for the AJ complex proteins β-catenin and N-Cad at 6 h after irradiation. White arrowheads indicate breaks in the AJ belt. Yellow arrowheads indicate regions of apparent delamination. n = 5; scale bars: 100 μm, 10 μm.

To further investigate whether radiation induced an EMT-like mechanism, we analyzed protein levels of AJ components and markers of EMT using western blotting. At 6 h after irradiation, expression of several proteins, including ZO-1, E-Cad, β-catenin and Qki5 were significantly reduced whereas α-catenin and N-Cad were not changed (Figs. 7b, c). This very much resembled the expression pattern of these proteins in ZIKV-NS2A infected microcephalic brains^48^. Moreover, immunostaining for β-catenin and N-Cad demonstrated that the integrity of the AJ belt was disturbed in brains of irradiated mice leading to an apparent delamination of cells from the ventricular surface (Figs. 7d, S8).

In brain tumors (gliomas), a mechanism resembling EMT is responsible for therapy resistance and cancer recurrence. PMT is the shift of glioma stem-like cells (GSCs) of the proneural subtype to the more aggressive mesenchymal subtype^49^. PMT can be induced by radiation both in mice^24^ and humans^25^. Halliday et al. showed that irradiation of proneural glioma in mice induces a p53-dependent DDR as well as a STAT3- and CEBP-dependent PMT^24^. Interestingly, most of the highly induced genes in irradiated glioma are among the top genes induced in irradiated embryonic mouse brains, in particular after 2 h and decreasing over time (Fig. S9a). These genes represent the p53-mediated DDR. Besides this, our RNA-seq results (Fig. 7e, f) and a GSEA analysis (Fig. S4c, S9b) showed that after 12 h the expression of proneural GSC markers such as *Ascl1, Bcan, Gpr17* and *Ttyh1* was reduced while mesenchymal GSC markers like *Alox5, Casp1, Lgals3* and *Ltbp2* were upregulated. Hence, our results demonstrate that the transcriptional response to radiation is very similar between the embryonic mouse brain and proneural glioma in adult mice.

## Discussion

The embryonic brain is very sensitive to DNA damage, especially DSBs. The occurrence of excessive DSBs during embryonic development is therefore a major cause of neurodevelopmental diseases, which often display microcephaly^20,21^. The main underlying mechanism, as in many other developmental syndromes associated with microcephaly^5^, is (hyper)activation of p53 leading to apoptosis of NPCs and a depletion of the neuronal progenitor pool. In this study, acute DNA damage induced by X-irradiation of mouse embryos at the start of neurogenesis leads to microcephaly resulting from a combination of apoptosis and p53-dependent premature neuronal differentiation.

The induction of ectopic neurogenesis was evident from a reduction in the number of Pax6+ RGCs, and the presence of Dcx+ and Tbr1+ immature neurons in the VZ of irradiated mice, coinciding with an increase in asymmetrically dividing RGCs. Here, we found that both apoptosis and premature neuronal differentiation were prevented by genetic inactivation of *Trp53* in the dorsal forebrain, resulting in a less severe microcephalic phenotype. The role of p53 in radiation-induced premature neuronal differentiation was furthermore supported by a functional genomics screen. We suggest that genes belonging to the *IR_unique* gene set are responsible for radiation-induced neuronal differentiation, which is strengthened by the fact that this gene set is also highly enriched during normal brain development and in several models of differentiating ESCs. Potential candidate genes are, among others, *Bloc1s2* and *Fbxw7*, both of which activate neuronal differentiation of NPCs by inhibition of Notch1 signaling^50,51^; *Btg2/Tis21*, which is a pro-differentiation gene that induces premature onset of consumptive divisions of NSCs and microcephaly^52^; *Baiap2/IRSp53* which promotes filopodia and neurite formation^53^; *Nexmif/Kidlia*, an X-linked intellectual disability gene which regulates neurite outgrowth^54^; *Nanos1*, an RNA-binding protein which promotes neurogenesis^55^, and *Arap2*, of which the human ortholog is one of 64 genes enriched in human outer radial glia^56^. Additionally, this gene set comprises currently uncharacterized p53 target genes of which some, like *D630023F18Rik (C2orf80* in humans) and *Sec14l5*, are highly upregulated during normal mouse brain development and neuronal maturation^31^. Notably, these genes may contribute to DNA damage-induced neuronal differentiation and therefore represent important targets for functional characterization.

Radiation-induced neuronal differentiation has been demonstrated *in vitro*^37,38,57,58^ although the underlying mechanisms remained largely unclear. To our knowledge, this is the first study demonstrating the activation of a p53-dependent transcriptional program as a mechanism to induce neuronal differentiation *in vivo*, although a role for p53 in cellular differentiation *in vivo* has been previously proposed in mammary stem cells, the airway epithelium and cancer stem cells^6^. Recently, it was also shown that high doses of radiation could drive ATM-dependent differentiation of neuroblasts in the adult SVZ^59^. Since ATM is the major activator of p53 in response to radiation-induced DSBs, it is also possible that this apparent ATM-dependence in fact reflects the effect of p53.

While the ultimate fate of the prematurely differentiated cells is unclear (e.g. functional integration, senescence, death), an important question is what underlies the cell’s decision to undergo either apoptosis or differentiation. Based on our results, it is tempting to speculate that the induction of apoptosis and differentiation gene signatures does not occur in the same cells. Transcriptomic analysis at the single cell level may answer this. Even then, the question remains what drives p53 to activate either of these signatures. Different possible explanations exist. For instance, the level of activation of p53 and its targets, the duration of their expression and the cellular context, irrespective of the amount of damage, may determine whether a cell undergoes cell cycle arrest or apoptosis^60^. Indeed, specific sets of downstream target genes which influence cell fate may be activated in a cell-specific manner depending on (i) the dynamic behavior of p53 expression^61^, (ii) posttranslational modifications of the p53 protein itself^62^ -mutant mice mimicking constitutive p53 acetylation have neuronal apoptosis and microcephaly^63^-, or (iii) the activation of specific splice variants of p53 explaining some of its pleiotropic activities^64^. It is, however, also possible that other cell-autonomous mechanisms, such as differences in their signaling landscape modulate the cell-specific choice between different p53-dependent transcription programs^4^. Another important question that remains to be answered is the extent to which radiation-induced premature differentiation contributes to the reduction in brain size. The fraction of prematurely differentiating cells seems lower than that of the apoptotic ones. One may therefore conclude that its contribution is smaller. However, if premature differentiation of an RGC would prevent it from subsequent rounds of division, it may still have a considerable impact. Also, if prematurely differentiated cells ultimately undergo apoptosis, then part of the observed apoptotic cells was prematurely differentiated in the first place. If possible, it would be interesting to generate a model in which only premature differentiation, but not apoptosis could be prevented in order to calculate the respective contributions of each of these cell fates to the phenotype.

The lack of an obvious brain phenotype in p53 cKO mice supports the idea that p53 is essentially dispensable for proper brain development *per se*, although it has been reported that a subset of p53-deficient animals develop exencephaly, a neural tube closure defect^65,66^. This discrepancy may be explained by the fact that in the *Emx1-Cre* model recombination only occurs at E9.5, when neural tube closure has already terminated. Thus, ablation of p53 at the stage of corticogenesis does not impact brain size and correct layering of the neocortex. Another explanation may be that although p53 expression levels are high during early brain development, its activation is very strictly controlled in the absence of cellular stress such that p53 activity is inherently low anyway^67^.

During neurogenesis, RGCs have an apicobasal polarity and delaminate from the apical membrane by downregulation of AJ complex proteins^22^. This process shares features with that of EMT and it has been shown that an EMT-like process precedes delamination of differentiating RGC daughter cells and their radial migration^46^. We found that in irradiated mice, several classical genes involved in EMT are induced. Also, AJ complex proteins such as E-Cad, ZO-1 and β-catenin were downregulated and disruption of apical AJs could be observed. This is very similar to the reduction in ZO-1 and β-catenin, but not N-Cad seen in ZIKV-NS2A infected embryonic mouse brains, displaying premature differentiation of RGCs^48^. The exact mechanism responsible for this EMT-like process still remains elusive. One possibility is that the AJ disruption is a secondary effect of apoptosis of nearby cells in the ventricular zone. During normal embryonic development, apoptosis is very important for correct morphogenesis and apoptotic cells can exert mechanical forces on the surrounding tissue which may affect cell-cell adhesion^68^.

The transcriptional response of the embryonic mouse brain to radiation very much resembled that of irradiated mouse glioma which was characterized by a p53-dependent apoptotic response and a p53-independent EMT-like mechanism driving a shift from proneural to mesenchymal cells^24^. In recent years, it has been postulated that the GSCs that confer glioma treatment resistance and tumor recurrence may arise from NSCs in the adult SVZ^69^. Importantly, two new studies showed that glioblastomas –high-grade gliomas-contain (outer) radial-glia-like cells^70,71^ which were hypothesized to be the cells of origin for glioblastoma development. Our observation of the pronounced similarities between the radiation response of the embryonic mouse brain and that of gliomas, supports this hypothesis. It furthermore indicates that the developing mouse brain may be used as a proxy to investigate certain aspects of glioma development and treatment response.

In summary, we have identified a novel *in vivo* role for p53 in activating neuronal differentiation via transcriptional regulation of differentiation-related genes in response to radiation-induced DNA damage. Further investigation of the specific functions of some of these genes in brain development under normal and genotoxic conditions and the mechanism responsible for the activation of a p53-dependent differentiation program are of pivotal importance to better understand p53-associated syndromes and to propose potential preventive strategies.

## Materials and methods

### Mouse husbandry

All animal experiments were handled in accordance with the Ethical Committee Animal Studies of the Medanex Clinic (EC_MxCl_2014_036). All of the animal experiments were carried out in compliance with the Belgian laboratory animal legislation and the European Communities Council Directive of 22 September 2010 (2010/63/EU). C57BL/6J mice were purchased from Janvier (Bio Services, Uden, The Netherlands). *Trp53* brain conditional knock-out (cKO) mice were obtained by breeding *Trp53^fl/fl^* mice (The Jackson Laboratory, Stock No. 008462) to *Emx1-Cre* mice (The Jackson Laboratory, Stock No. 005628). Mutants were genotyped by PCR. All the mice were housed under standard laboratory conditions with 12-h light/dark cycle. Food and water were available *ad libitum*. Female and male mice were coupled during a 2-h time period in the morning, at the start of the light phase (07:30 to 09:30 am) in order to ensure synchronous timing of embryonic development. The morning of coupling was considered embryonic day 0 (E0).

### Cell culture

For culturing adherent primary mouse NPCs fetal brains were dissected from E15 mouse fetuses and prefrontal cortices were separated. These were dissociated by gentle pipetting in Accutase (A6964, Sigma) and NPCs were cultured as monolayers in proliferation medium consisting of Dulbecco’s Modified Eagle Medium (DMEM)/F-12 with Glutamax (31331-093, Gibco) supplemented with 1x B-27 (17504-044, Gibco), 1x N-2 supplement (17502-048, Gibco), 10 ng/ml of recombinant murine Epidermal Growth Factor (EGF, 315-09, Peprotech) and 20 ng/ml recombinant human Fibroblast Growth Factor-basic (FGF-2, 100-18B, Peprotech) onto poly-Lysine D coated cell culture plates (Corning). Cells were subcultured every 3 to 4 days when they reached 70-80% confluence.

Neuro-2a cells were cultured in DMEM (61965-026, Gibco) supplemented with 10% fetal bovine serum (FBS, 10270-106, Gibco) and 1x non-essential amino acids (NEAA, 11140-035, Gibco) (proliferation medium) at 37°C with 5% CO2. For differentiation experiments, cells were subcultured in differentiation medium consisting of DMEM with 1% FBS (Gibco) and 1x NEAA (Gibco) supplemented with 10 μM of freshly added retinoic acid (R2625, Sigma). For experiments using the p53 transcriptional inhibitor α-pifithrin (α-PFT; P4236, Sigma-Aldrich, Diegem, Belgium), cells were treated with either 10 mM α-PFT or 1% DMSO 90 min prior to irradiation.

### Irradiation procedures

For irradiation experiments, pregnant dams at E11 or E14 were given a single dose of whole-body radiation (1 Gy), by using an X-Strahl 320 kV (0.13 Gy/min, inherent filtration: 3 mm of Be, additional filtration: 3.8 mmAl + 1.4 mm Cu + DAP, tube voltage: 250 kV, tube current: 12 mA, sample distance: 100 cm, beam orientation: vertical) in accordance to ISO 4037. Control mice were taken as well to the radiation facility but were not placed within the radiation field (sham-irradiation). For all experiments, embryos from at least two different litters were used as biological replicates.

Irradiation of NPCs (1 Gy) was performed using the same settings as for mice, for Neuro-2a cells (8 Gy) a dose-rate of 0.5 Gy/min was used.

### Brain size assessment

Brain size measurements were performed at postnatal day 1 (P1). The brains of the pups exposed to radiation *in utero* at E11 or E14 were isolated and imaged on a Leica stereo dissection microscope. The surface area of both cortices was measured and analyzed using ImageJ.

### Immunohistochemistry and imaging

Mouse fetuses (E11) or brains of E15 fetuses were harvested and fixed in 4% paraformaldehyde (PFA) dissolved in phosphate buffered saline (PBS) at 4°C overnight. The samples were then washed three times (5 min each) with PBS and stored at 4°C in 70 % ethanol till further manipulations. Next, the samples were processed with the Excelsior^™^ AS Tissue Processor (Thermo Scientific) and embedded using the paraffin-based embedding station (Ventana Discovery Ultra, Roche). Following the embedding, the brains were cut into 7-μm thick coronal sections on a microtome (Microm HM 340 E, Thermo Scientific), and mounted on SuperFrost^™^ Plus glass slides (Thermo Scientific).

P2 pups were transcardially perfused with a 0.9% NaCl solution and fixed in 4% PFA at 4°C overnight. After washing, samples were cryopreserved through a 10%-20%-30% series of sucrose dissolved in PBS, and embedded in frozen section medium (Richard-Allan Scientific^™^ Neg-50^™^). P2 brains were cut into 10-μm thick coronal cryosections (Cryostar NX50, Thermo Scientific) and mounted on SuperFrost^™^ Plus glass slides.

Before staining, paraffin sections were deparaffinized in xylene, rehydrated in graded solutions of ethanol, and boiled in citrate buffer, pH6 (Dako) for antigen retrieval. Next, the sections were incubated three times (5 min each) in permeabilization solution (0.3 % Triton X-100) before blocking for 2 h either with Normal Goat Serum (Invitrogen) in Tris-NaCl blocking buffer (1:5) or PBS containing 1 % BSA (Sigma Life Science) and 0.3 % Triton X 100 (Sigma Life Science). After blocking, the sections were then incubated overnight at 4°C in blocking solution containing the primary antibody. The following antibodies at the indicated dilution were used: anti-p53 (mouse, 1:100, Santa Cruz Biotechnology (sc-126)) anti-53BP1 (rabbit, 1:100, Novus Biologicals (NB 100-304ss)), anti-PH3ser10, (rabbit, 1:1000, Cell Signaling (3377)), anti-Pax6 (rabbit, 1:200, Biolegend (901301)), anti-DCX (Goat, 1:500, Santa Cruz Biotechnology (sc-8066)), anti-cleaved Caspase-3 (rabbit, 1:100, Biovision (3015-100)), anti-BrdU (rat, 1:300, Bio-Rad (BU1/75)), anti-Ctip2 (rat, 1:500, Abcam (ab 18465)), anti-Satb2 (mouse, 1/100, Abcam (ab51502)), anti-Tbr1 (rabbit, 1:100, Abcam (ab31940), anti-Tbr2 (rabbit, 1:100, Abcam (ab23345), anti-γ-tubulin (goat, 1:100, Santa Cruz Biotechnology (sc-7396)), anti-β-catenin (mouse, 1:100, Santa Cruz Biotechnology (sc-7963)), anti-N-cadherin (rabbit, 1:350, Abcam (ab18203)), anti-Tuj1 (mouse, 1:1000, Sigma Life Science (T5076)).The following day, sections were washed three times (5 min each) with tris-buffered saline, 0.1% Tween20 and incubated 2 h at room temperature with the appropriate Alexa Fluor-405, −488 or −568 (Invitrogen) secondary antibody diluted 1:200 in blocking solution. For CC3 staining secondary antibody was followed by signal amplification using TSA Plus Cyanine 3 System (PerkinElmer). Following the incubation with the secondary antibody, sections were washed three times (5 min each) and counterstained for nuclei, with 4’6-diamidiono-2-phenylindole (DAPI, Sigma-Aldrich) for 15 min. Finally, the slides were mounted with mowiol and images were taken using 20x or 40x air objectives on a Nikon Eclipse Ti-E inverted microscope.

### EdU/BrdU cumulative pulse-labeling experiments

Sequential administration of the thymidine analogues, 5-ethynyl-2’-deoxyuridine (EdU) (Click-iT Plus EdU Kit, Invitrogen) and 5-bromo-2’-deoxyuridine (BrdU) (Sigma-Aldrich) was used for cell cycle length assessment. Briefly, pregnant dams (E11) were given intraperitoneal injections of EdU (10 mg/kg of body weight), and BrdU (50 mg/kg of body weight), 22 h and 23.5 h respectively after irradiation. Half an hour later fetuses were harvested and processed as mentioned above. EdU detection was done using the Click-iT Plus EdU Alexa Fluor 488 Imaging Kit (Invitrogen) according to the manufacturer’s protocol. Subsequently, BrdU (Alexa 568) and Pax6 (Alexa 405) stainings were performed as indicated above. The total cell cycle length and duration of S-phase of Pax6 positive NPCs were calculated as follows: the interval during which cells can incorporate EdU, but not BrdU (T_i_) is 1.5 h. The total number of Pax6+ cells (cycling fraction), the number of Pax6+ cells in S-phase (S fraction, Pax6+ EdU+ BrdU+) and the number of Pax6+ cells in the leaving fraction (L fraction, Pax6+ EdU+ BrdU-) were analyzed using ImageJ. Duration of the S-phase (T_s_) was then calculated using the following equations^72^:

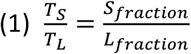

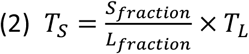

Because the ratio of the length of any one period of the cell cycle to that of another period is equal to the ratio of the number of cells in the first period to the number in the second period, the ratio between the number of cells in the S fraction and the L fraction is equal to the ratio between T_S_ and T_i_ (= 1.5 h), and therefore T_L_ = T_i_.

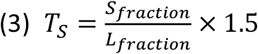

This could be used to calculate the total cell cycle length (T_C_) was calculated using:

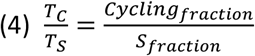

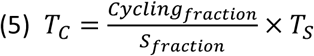

### RNA library preparation for RNA sequencing (RNA-Seq)

For RNA-seq, pregnant dams were (sham-)irradiated at E11 after which they were sacrificed for dissection of fetuses after 2 h, 6 h and 12 h. For each condition 3 fetuses were used from at least 2 different mothers. Fetal cortices were dissected and stored in RLT Plus lysis buffer (Qiagen) at −80°C until RNA extraction using the RNeasy Mini Kit (Qiagen). RNA quality was determined using the RNA nano assay on a 2100 Bioanalyzer (Agilent Technologies). All samples had RNA Integrity Numbers >9.10. Triplicate RNA-Seq libraries were prepared according to the TruSeq stranded mRNA protocol (Illumina). Briefly, 200 ng of total RNA was purified using poly-T oligo-attached magnetic beads to end up with poly-A containing mRNA. The poly-A tailed mRNA was fragmented and cDNA was synthesized using SuperScript II and random primers in the presence of Actinomycin D. cDNA fragments were end repaired, purified with AMPure XP beads, A-tailed using Klenow exo-enzyme in the presence of dATP. Paired end adapters with dual index (Illumina) were ligated to the A-tailed cDNA fragments and purified using AMPure XP beads. The resulting adapter-modified cDNA fragments were enriched by PCR using Phusion polymerase as followed: 30 s at 98°C; 15 cycles of 10 s at 98°C, 30 s at 60°C, 30 s at 72°C; 5 min at 72°C. PCR products were purified using AMPure XP beads and eluted in 30 μl of resuspension buffer. One microliter was loaded on an Agilent Technologies 2100 Bioanalyzer using a DNA 1000 assay to determine the library concentration and for quality check.

### Bridge amplification, sequencing by synthesis and data analysis

Cluster generation was performed according to the TruSeq SR Rapid Cluster kit v2 (cBot) Reagents Preparation Guide (Illumina). Briefly, 36 RNA-Seq libraries were pooled together to get a stock of 10 nM. One microliter of the 10 nM stock was denaturated with NaOH, diluted to 6 pM and hybridized onto the flowcell. The hybridized products were sequentially amplified, linearized and end-blocked according to the Illumina Single Read Multiplex Sequencing user guide. After hybridization of the sequencing primer, sequencing-by-synthesis was performed using the Illumina HiSeq 2500 with a single read 50-cycle protocol followed by dual index sequencing, producing ~782 million 50-bp reads (~22 million reads per condition). Reads were aligned against the GRCm38 genome using HiSat2 (version 2.0.4)^73^. Counts were generated for each gene from the Ensembl transcriptome analysis of GRCm38, using htseq-count (version 0.6.0)^74^. Genes with an FDR adjusted *P* < 0.05 were considered as differentially expressed.

Gene Ontology analysis (Figs. 3e, 6c, 6e, 6g) was performed using the *Investigate Gene Sets* tool from the MSigDB (v7.0) using GO Biological Process as the reference with FDR q-value < 0.05. GSEA analysis^35^ was performed with a weighted enrichment statistic and a signal-to-noise metric for gene ranking. GSEA for Figs. 3d and S3, was performed with MSigDB Hallmark gene sets and curated gene sets, respectively, based on RNA-seq results. GSEA for Figs. 6h and 6i was performed with microarray results from embryonic brain development^31^ and DNA damage-induced ESC differentiation^75^, respectively. Transcription factor enrichment analysis to identify transcriptional regulators was performed using Enrichr^76,77^. Genes significantly upregulated (FDR < 0.05) in WT animals were visualized as a network (Fig. 3b) using Cytoscape (v3.6.1)^78^. Edges between p53 and other genes were based upon Enrichr analysis. RNA sequence data reported in this paper have been deposited in the Gene Expression Omnibus (GEO) database (GSE140464).

### Quantitative reverse transcriptase PCR (qRT-PCR)

RNA was extracted from cells using the RNEasy Mini kit (Qiagen) and eluted in 30 μl of RNase-free water. This was used for cDNA synthesis using the GO-Script Reverse Transcriptase kit (Promega) using 1 μl of random hexamer primers and 3.75 mM MgCl2 per 20-μl reaction. Quantitative PCR was then performed using an ABI7500 Fast instrument and the MESA Green qPCR MasterMix Plus for SYBR assay (Eurogentec). Relative expression was calculated via the Pfaffl method^79^ using *Gapdh* and *Polr2a* as references gene. Primers used for qRT-PCR are listed in table S7. For all qRT-PCR experiments the specificity of the primers was validated using a melting curve.

### MicroRNA microarrays and data analysis

Total RNA, including miRNA was isolated from brains of E11 mouse fetuses (3 biological replicates) at 2 h after irradiation using the miRNeasy Mini Kit (Qiagen). RNA was subsequently processed for hybridization to GeneChip miRNA 4.0 microarrays (Affymetrix) using the FlashTag^™^ Biotin HSR RNA Labeling Kit (Affymetrix) according to the manufacturer’s instructions. Briefly, 1000 ng of total RNA was used for poly (A) tailing, followed by biotin labeling using the FlashTag^™^ Biotin HSR Ligation Mix. Biotin-labeled microRNAs were then hybridized to microarrays at 48°C for 16 h. Microarrays were washed and stained with streptavidin at 35°C in a Fluidics Station 450 (Affymetrix) and scanned with a GeneChip Scanner 7G (Affymetrix).

Microarray data analysis was performed using Partek Genomcs Suite (v7.17.1018). CEL-files were imported using a customized Robust Multi-array Average algorithm (background correction for probe sequence, quantile normalization, log2 transformation of intensity signals). Mouse microRNAs with a *P* <0.01 (Student’s *t*-test) were considered as differentially expressed.

### Western blotting

Equal amounts of E11 brain homogenates were loaded onto 4-15% TGX precast gels (BioRad) and transferred to nitrocellulose membranes (or polyvinylidene difluoride for alpha-catenin detection). Membranes were incubated overnight with antibodies directed to ZO-1 (61-7300, ThermoFisher Scientific, 1/100), E-cadherin (ab76055, Abcam, 1/200), beta-catenin (sc7963, Santa Cruz Biotechnology, 1/2500), alpha-catenin (C2081, Sigma-Aldrich, 1/5000), N-cadherin (ab18203, Abcam, 1/1000) and QKI-5 (A300-183A, Bethyl Laboratories, 1/5000), followed by 45 min incubation with the appropriate HRP-conjugated antibodies (Invitrogen). Bands were visualized using the luminol-based enhanced chemiluminescent kit (ClarityTM Western ECL Substrate, BioRad) and a Serva Purple total protein stain (SERVA Electrophoresis GmbH) was used for protein normalization. Blots were imaged using a Fusion FX (Vilber Lourmat) imaging system and band intensities were measured with the FusionCapt Advance and Bio1D software packages (Vilber Lourmat) for semi-quantitative analysis.

### Live cell imaging

For live cell imaging, mouse NPCs and Neuro-2a cells (8000 cells/well) were seeded in 96-well plates and placed in an IncuCyte ZOOM (Essen Bioscience) immediately after subculturing. Phase-contrast images were captured with a 10x (Neuro-2a) or 20x (NPCs) objective, every 2 h for a total of 72 h to 96 h. Per well, two images were captured for every time point and per experimental condition 4-6 wells were used as technical replicates. All live cell imaging experiments were replicated independently.

### Statistical analysis

Statistical analysis was performed using GraphPad Prism 7.02. Comparisons between control and irradiated C57BL/6 mice were performed using unpaired two-tailed Student’s *t*-test. Comparisons between control and irradiated WT and cKO mice were performed using one-way ANOVA. A *p*-value of <0.05 was considered statistically significant. More details about statistical tests and numbers of replicates are indicated in figure legends.

## Acknowledgements

The authors wish to thank Ann Janssen, Amelie Coolkens, Annick Francis, Mieke Neefs, Lisa Daenen, Brit Proesmans, and Bent Hubrecht for excellent technical assistance. This work was supported by the 7^th^ European Framework Programme (CEREBRAD - GA: 295552 to M.A.B.), the Fonds Wetenschappelijk Onderzoek (G0A3116N to D.H. and R.Q.) and the Aspirant Wetenschappelijk Medewerker programme of the Belgian Nuclear Research Centre (to A.C.M.M., M.V., and T.V.).

## Author contributions

R.Q., M.A.B., D.H., L.M. conceived the study. A.C.M.M., R.Q., M.V., T.V., H.B.F. designed and performed the experiments. R.Q., A.C.M.M., M.V., M.M., W.F.J.V.I., T.V., H.B.F. analyzed the data. W.H.D.V. developed software. D.H., R.Q., M.A.B., S.B. acquired funding. R.Q. wrote the draft manuscript. A.C.M.M., M.V., T.V., M.M., W.F.J.V.I., W.H.D.V., L.M., M.A.B., D.H. reviewed and edited the manuscript.

## Competing interests

The authors declare no competing interests.

## Supplemental figure legends

**Figure S1:**
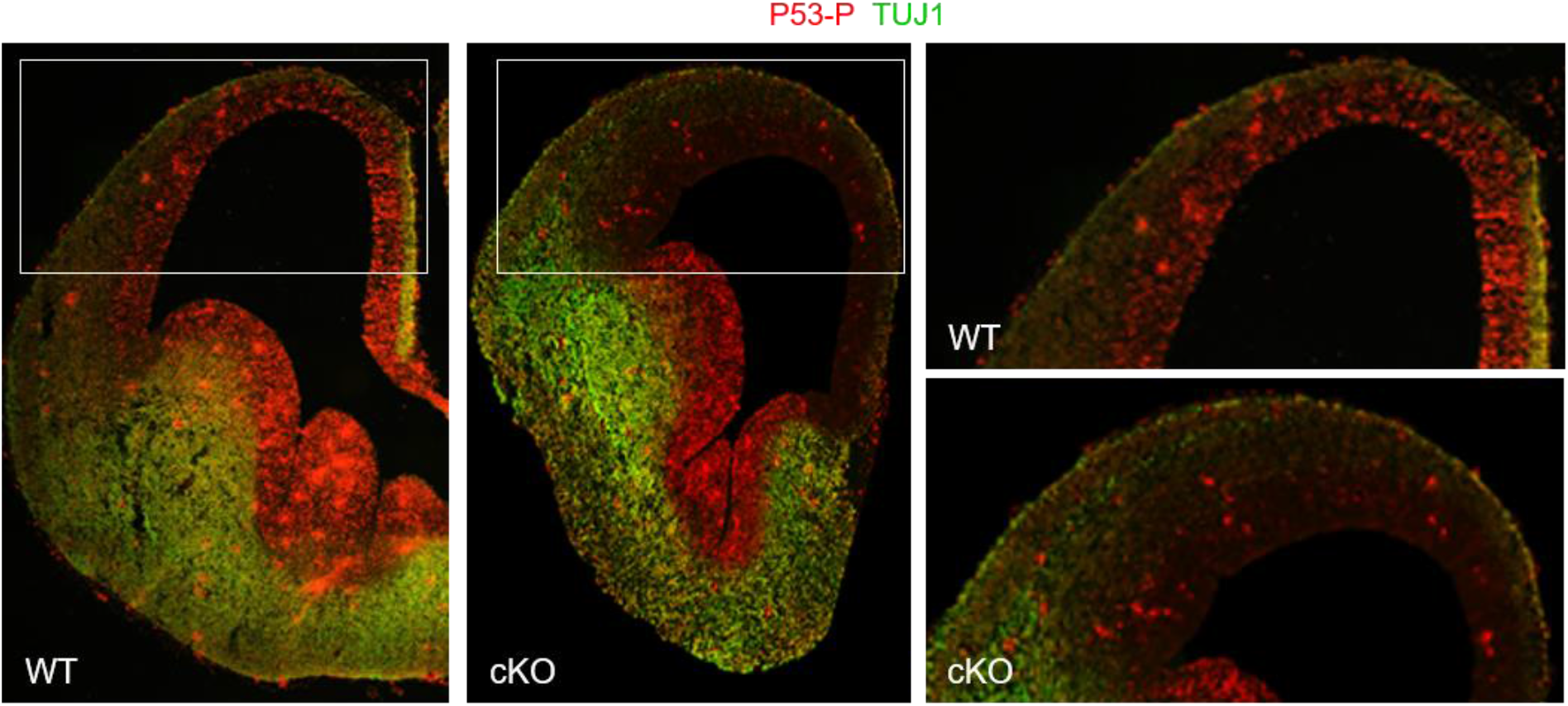
Specific ablation of phospho-p53 expression in the dorsal neocortex at E14. Pregnant mice were irradiated at E14 and embryos were dissected 2 h later for immunostaining for phospho-p53 and TUJ1.

**Figure S2:**
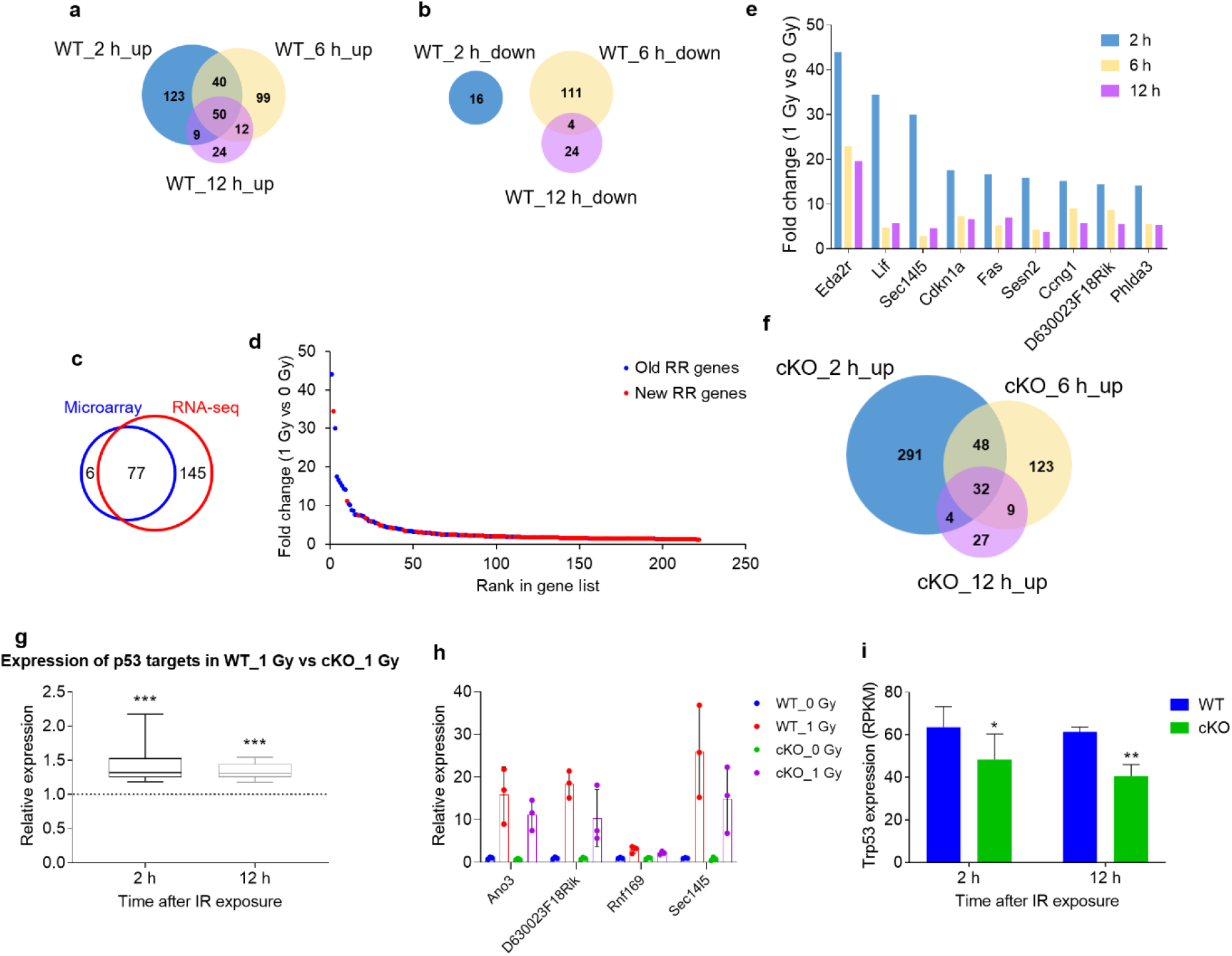
Radiation-induced changes in gene expression. (**a, b**) Venn-diagrams showing overlap in upregulated (**b**) and downregulated (**c**) genes in WT mice between the different time points. (**c**) Overlap between genes identified as upregulated after 2 h in WT mice by microarray and RNA-seq. RR: radiation-responsive. (**d**) Genes ranked according to their fold change in expression in irradiated versus control brain at 2 h after irradiation. (**e**) Time-dependent gene expression of the top 9 genes induced after 2 h. (**f**) Venn-diagram showing overlap in upregulated genes in cKO mice between the different time points. (**g**) Difference in fold change induction of p53 targets between WT and cKO mice at 2 h after irradiation. \*\*\**P* < 0.001 (Paired Student’s *t*-test). (**h**) qRT-PCR analysis of *Ano3, D630023F18Rik, Rnf169* and *Sec14l5* expression in WT and cKO cortices at 2 h after irradiation. (**i**) Trp53 mRNA expression in WT vs cKO mice after irradiation. n = 3; \**P* < 0.05, \*\**P* < 0.01 (Paired Student’s *t*-test).

**Figure S3:**
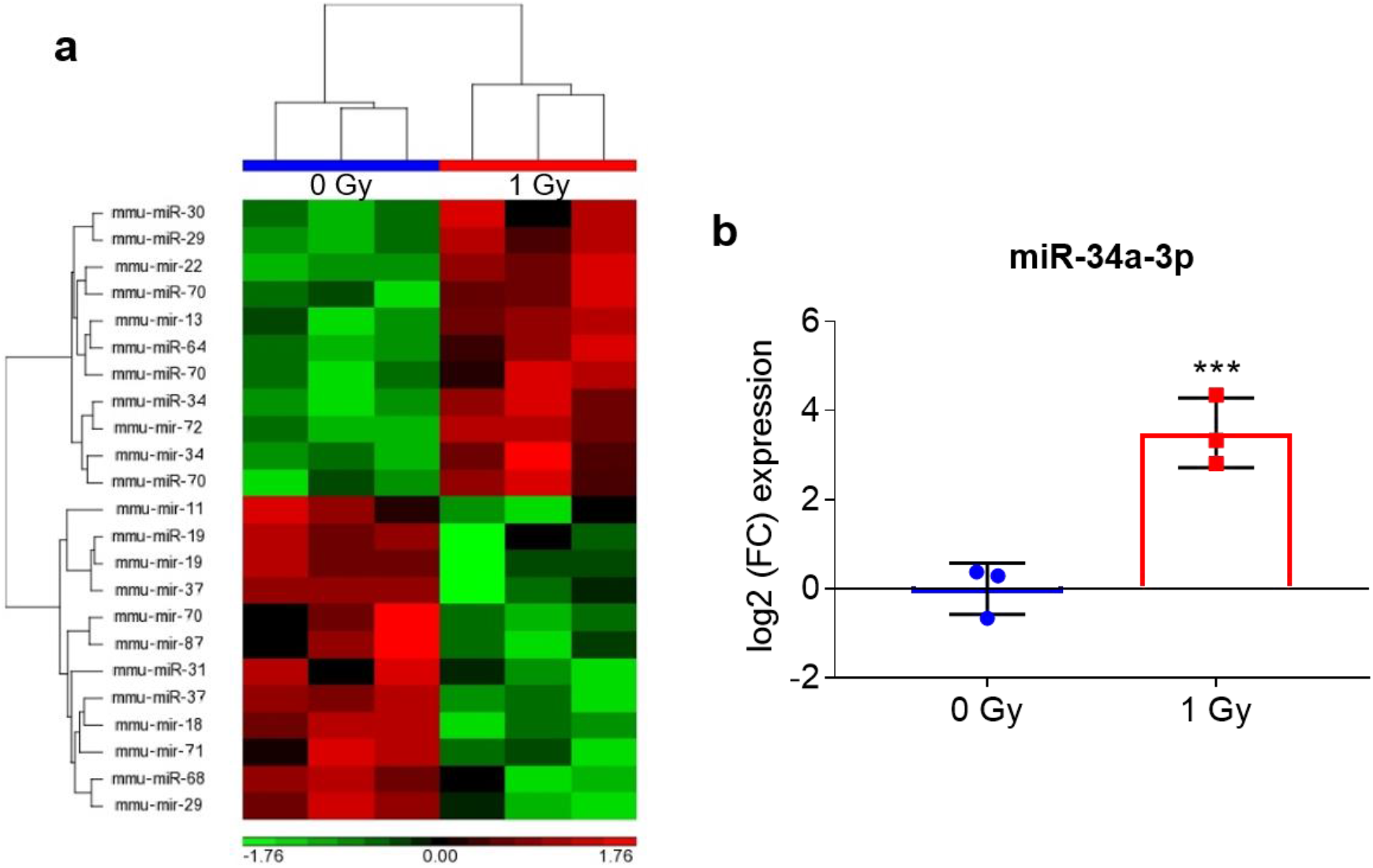
Radiation-induced changes in expression of microRNAs. (**a**) Heatmap depicting the most significantly changed microRNAs, of which the p53 target miR-34a (**b**) was the most extensively upregulated. \*\*\**P* <0.001 (Student’s *t*-test).

**Figure S4:**
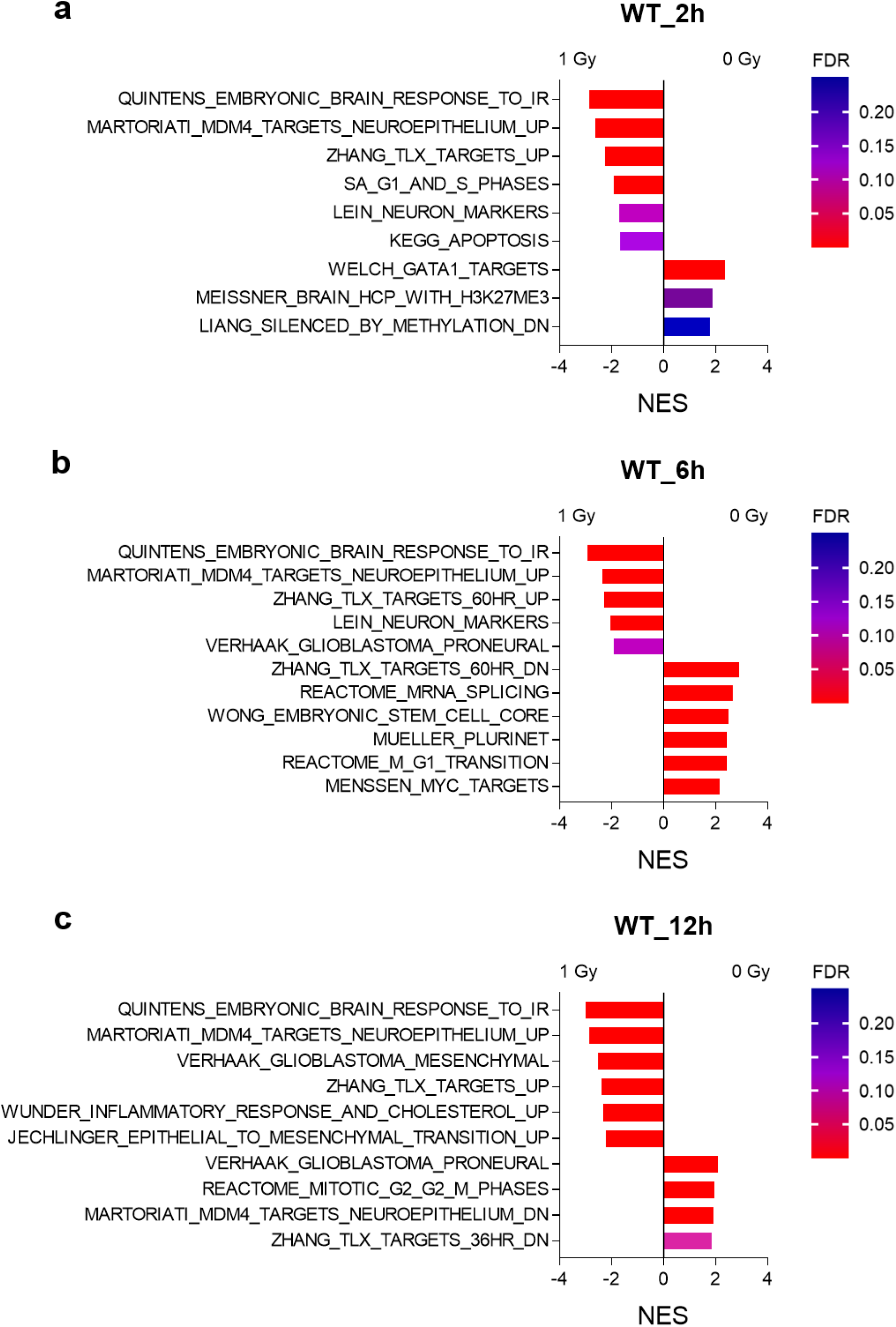
Gene Set Enrichment Analysis of gene expression profiles at 2 (**a**), 6 (**b**) and 12 h (**c**) post-irradiation. Selected gene sets are presented. The full lists can be found in Tables S2-S4.

**Figure S5:**
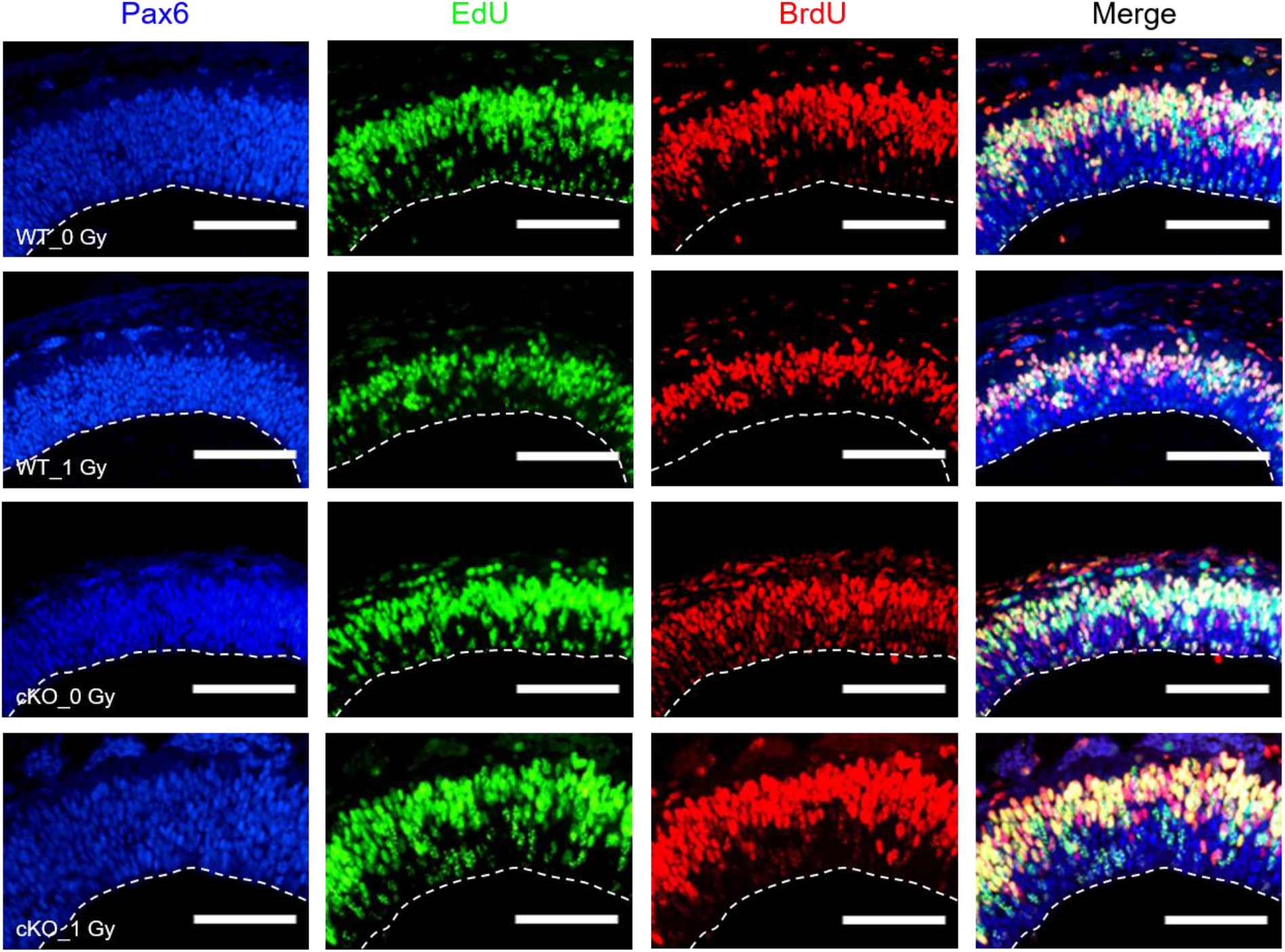
Cell cycle length of neocortical radial glia is not altered at 24 h after irradiation. To calculate cell cycle length, mouse fetuses underwent a sequential EdU, BrdU administration paradigm (see methods). Based on stainings for Pax6 (blue), EdU (green) and BrdU (red) both the duration of the total cell cycle and that of the S-phase could be calculated (see methods and Fig. 5g). n = 6 (WT_0 Gy) or n = 5 (all other conditions); scale bars: 100 μm.

**Figure S6:**
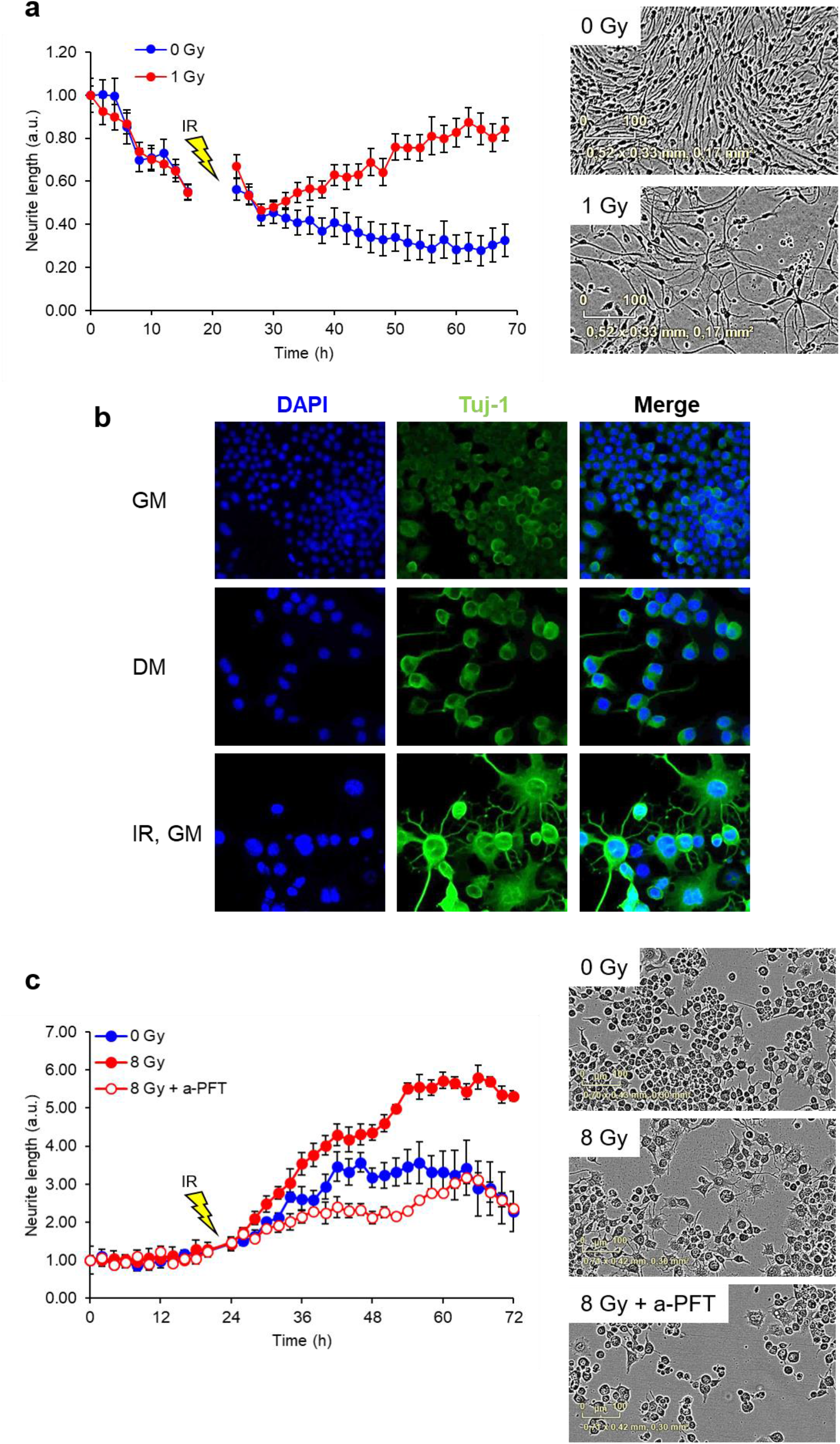
Exposure to radiation induces differentiation of mouse neural progenitor cells (NPCs) and Neuro-2a cells in a p53-dependent manner. (**a**) Mouse NPCs were irradiated (IR) at day-in-vitro (DIV) 2 and neurite outgrowth was analyzed via live cell imaging for two more days. (**b**) Neuro-2a cells at DIV 3 in growth medium (top), differentiation medium (middle) and growth medium after irradiation at DIV1 (bottom). Note the enlargement of the nuclei in differentiated cells and the multipolar neurite outgrowth in irradiated cells as compared to the uni-/bipolar cells in differentiation medium. (**c**) Neurite outgrowth analysis during live cell imaging of Neuro-2a cells shows enhanced neurite outgrowth in irradiated (IR) cells as compared to sham-irradiated cells which could be prevented by prior administration of the p53 inhibitor a-pifithrin (a-PFT). In all panels, data represent mean ± s.e.m. GM: proliferation medium; DM: differentiation medium.

**Figure S7:**
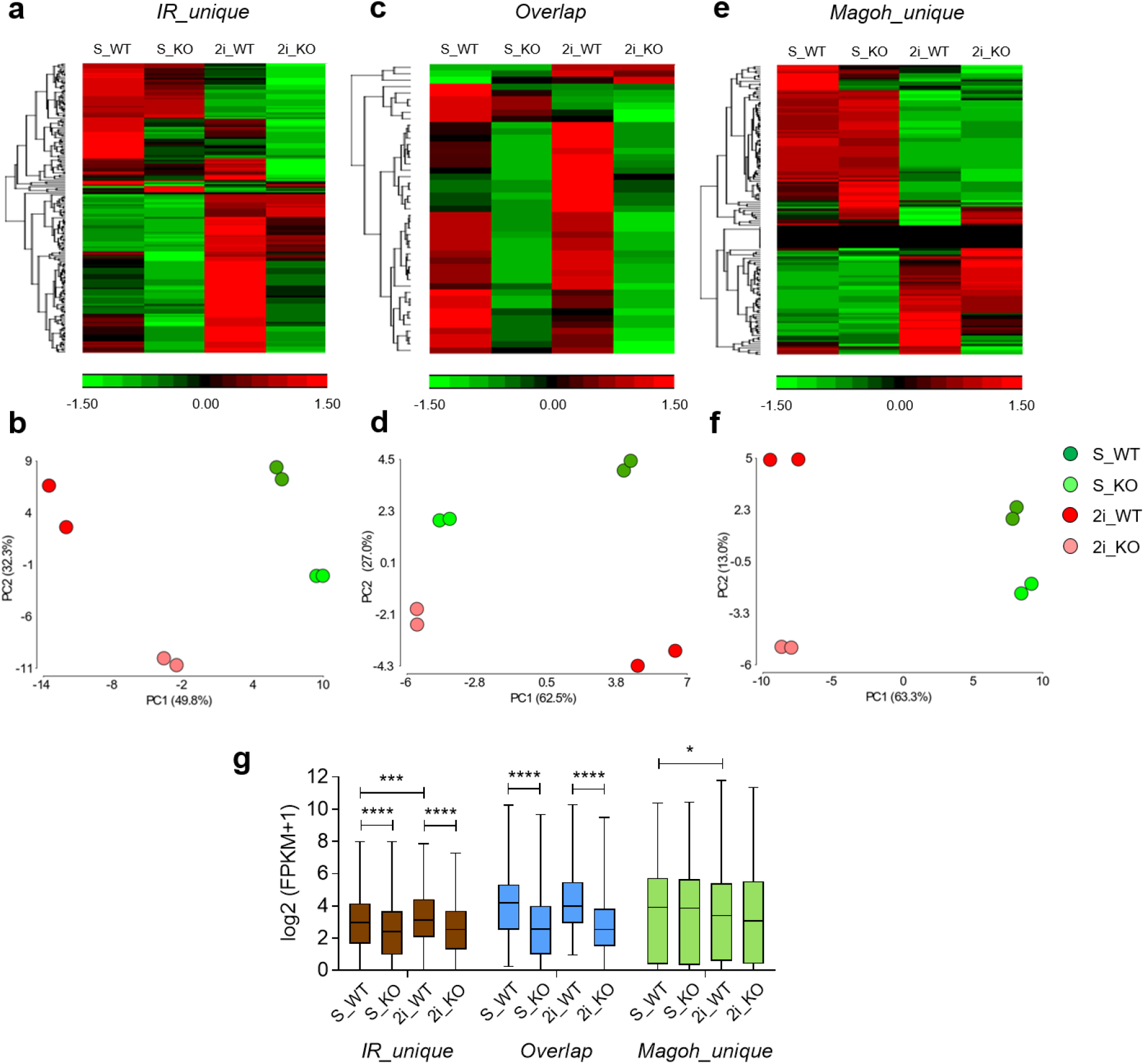
Expression profiles of *IR_unique, Overlap*, and *Magoh_unique* gene sets in wild-type compared to TP53^-/-^ (KO) ground state (2i) and serum-cultured (S) embryonic stem cells (data from^41^). (**a-c**) Heatmaps depicting gene expression profiles. (**b-f**) Principal component analysis indicates the variation explained by the respective gene sets. Please not that for *IR_unique* and *Magoh_unique* the cell culture conditions are in PC1 while for the *Overlap* genes PC1 represents the genotype. (**g**) RNA-seq counts of *IR_unique, Overlap*, and *Magoh_unique* genes. Data represent mean ± S.D. \**P* < 0.05, \*\*\**P* < 0.001, \*\*\*\**P* < 0.0001 (Two-way ANOVA).

**Figure S8:**
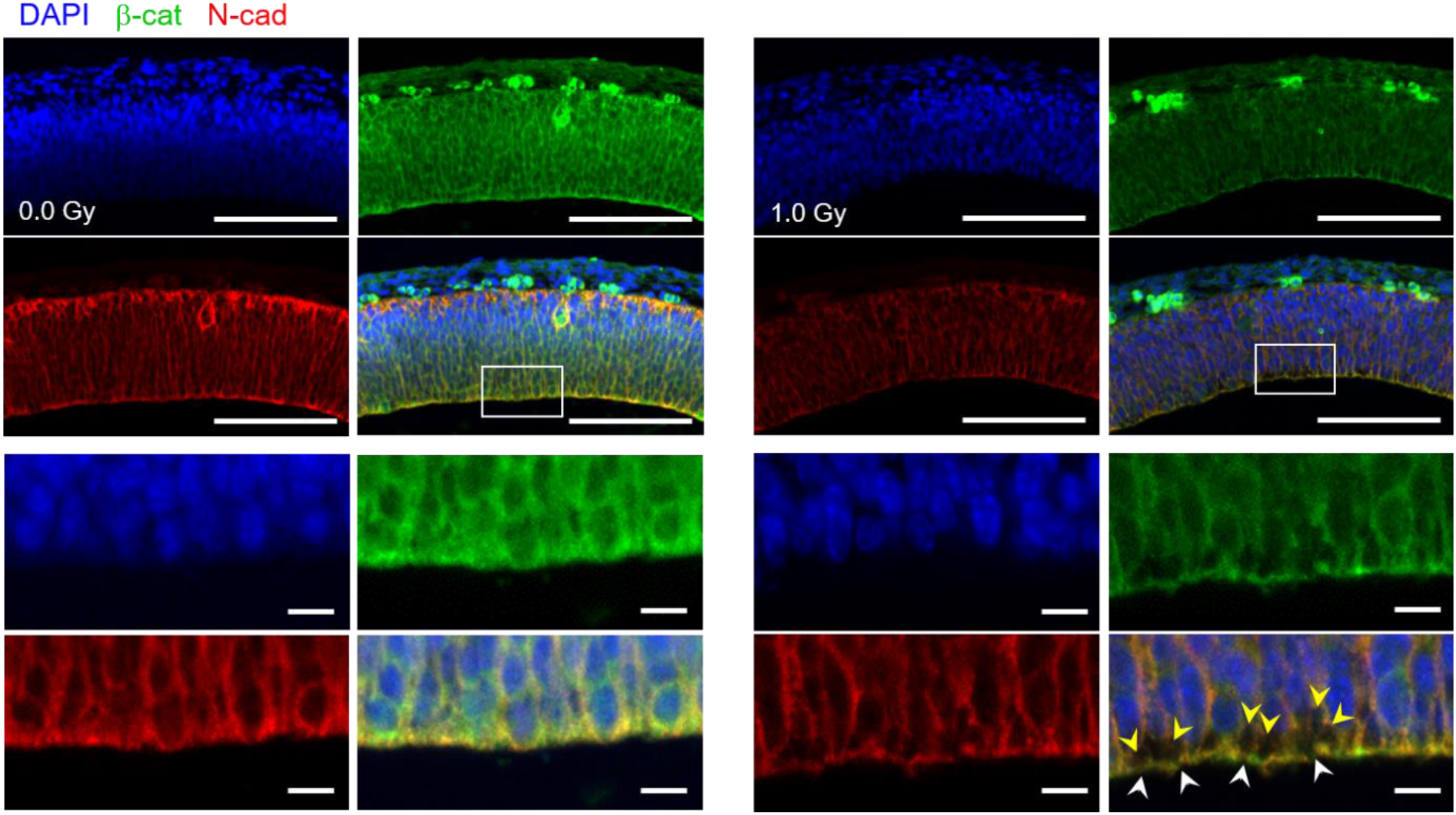
Disruption of the adherens junction (AJ) belt at the apical surface of the ventricular zone. Representative images of immunostaining for the AJ complex proteins β-catenin and N-Cad at 6 h after irradiation. White arrowheads indicate breaks in the AJ belt. Yellow arrowheads indicate regions of apparent delamination. n = 5; scale bars: 100 μm, 10 μm.

**Figure S9:**
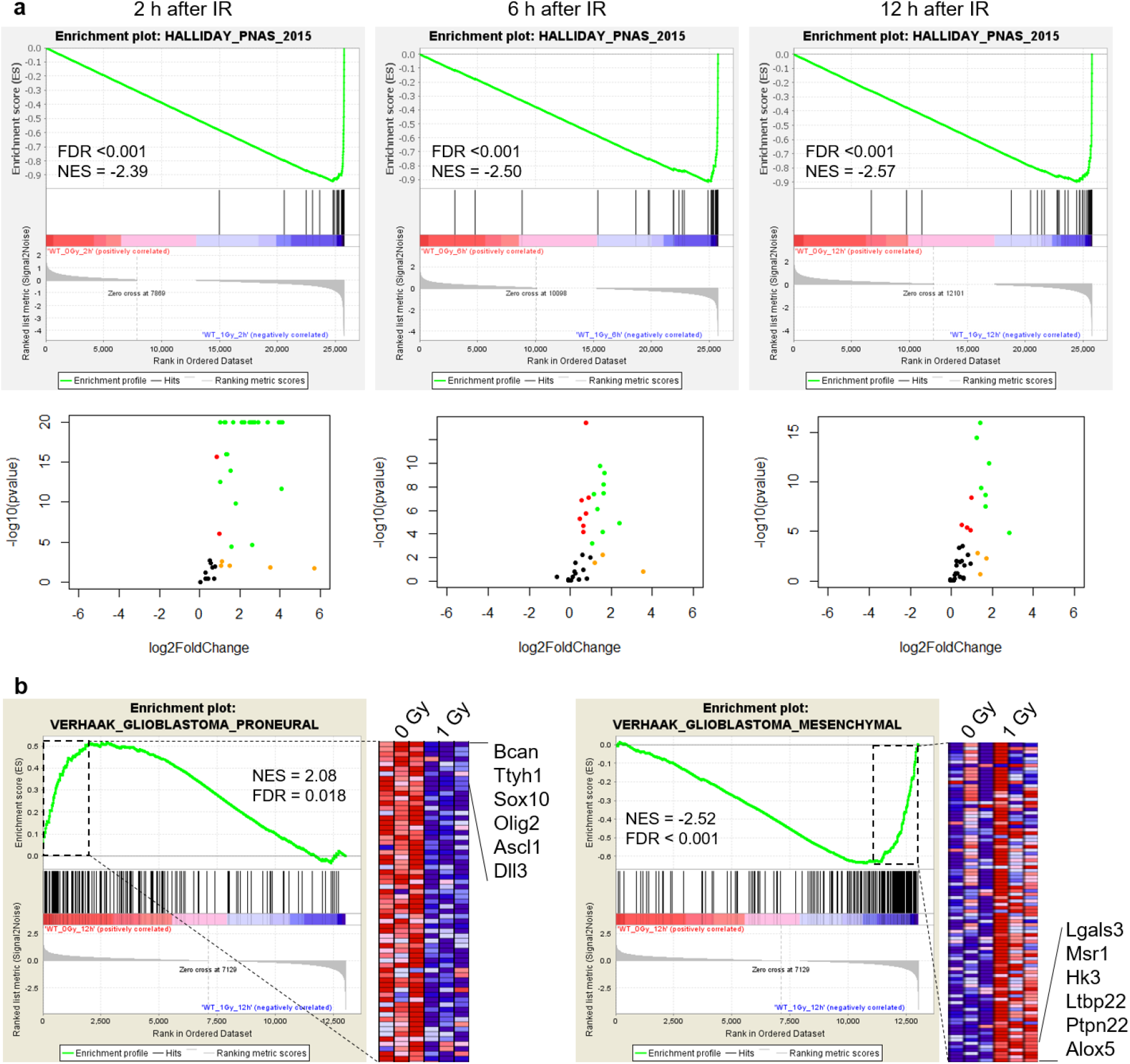
Irradiation of the embryonic mouse brain and proneural glioma activate a similar transcriptional response. (**a**) The top 42 genes induced by radiation in glioma (taken from^24^) were used to generate GSEA enrichment plots (upper panels) and volcano plots (lower panels) from embryonic brains at 2 h (left), 6 h (middle) and 12 h (right) following irradiation. Green points indicate genes with a FDR-corrected *p*-value <0.05 and a Log2 fold change >1; Red points indicate genes with a FDR-corrected *p*-value <0.05; Orange points indicate genes with a Log2 fold change >1. For genes with a *p*-value = 0, the *p*-value was arbitrarily set at 10^E-20^ to calculate the −Log10(*p*-value). (b) GSEA enrichment plots of proneural (left) and mesenchymal (right) gene signatures in embryonic brains at 12 h following irradiation.

## Notes

### Competing Interest Statement

The authors have declared no competing interest.

